# Stability-driven multi-omics integration for reproducible latent structure

**DOI:** 10.64898/2026.06.23.734064

**Authors:** Haibin Guan, Maaike van Gerwen, Seunghee Kim-Schulze, Elena Colicino, Georgia Dolios, Lauren M. Petrick

## Abstract

High-dimensional multi-omics data integration offers novel opportunities to characterize complex biological systems. Even though sampling variability frequently compromises findings, particularly in small cohorts, the reproducibility and generalizability of the derived latent structures are insufficiently evaluated. We propose a Stability-driven framework for multi-omics integration that combines sparse generalized canonical correlation analysis with repeated cross-validation, out-of-sample projection, and systematic evaluation of both component-level and feature-level stability. We apply this framework to untargeted metabolomic and Olink targeted inflammation proteomic profiles in a thyroid cancer case-control cohort (n = 162). Our Stability-driven integration identified reproducible metabolomic and proteomic latent components that showed consistent out-of-sample disease associations and tracked temporally structured changes relative to time to diagnosis. The proposed framework provides a generalizable strategy for identifying reproducible latent structures that improve robustness of biological inference in multi-omics studies.

## Introduction

The simultaneous measurement of multiple molecular layers—including genomics, transcriptomics, metabolomics and proteomics—within the same biological samples offers novel opportunities to investigate complex disease processes at a systems level ^1^. To extract a meaningful biological signal that represents shared variation across the omics layers, multi-omics integration tools have been widely developed^2^. These include covariance or correlation-based multiblock approaches such as Regularized Generalized Canonical Correlation Analysis (RGCCA) and Data Integration Analysis for Biomarker discovery using Latent cOmponents (DIABLO)^3^, which respectively extend canonical correlation analysis (CCA)^4^ or partial least squares (PLS) to multiple omics layers, as well as Bayesian factor analysis such as Multi-Omics Factor Analysis (MOFA) or MOFA+^5,6^, and matrix decomposition and factorization approaches (e.g., Joint and Individual Variation Explained [JIVE]^7^, scikit-fusion^8^ and Integrative Non-negative Matrix Factorization [intNMF]^9^). Unlike single-omics analyses that examine each omics layer independently, these multi-omics integration tools reduce the complexity of high-dimensional data by identifying a small number of latent components, with each sample receiving a score reflecting its overall coordinated variation across omics layers.

However, despite their increasing use, a critical challenge remains insufficiently addressed: whether the latent components—and the features that define them—are reproducible under sampling variability. In typical biomedical studies, the number of molecular features far exceeds the sample size^10–12^, and features within each omics layer are often highly correlated. Under these conditions, sparse multiblock models require extensive regularization^4^ and parameter tuning, which can lead to overfitting, instability in feature selection^12^, and sensitivity of latent components to sampling variability. In the fields of neuroimaging and high-dimensional regression, unstable latent component structures and associated feature weights are observed even at moderate sample sizes^11^, indicating possible reproducibility concerns of the inferred biological signals.

A key limitation of current practice is that stability is rarely evaluated explicitly. Most multi-omics studies rely on a single fitted model and interpret latent components and selected features based on in-sample associations. However, for dimensionality-reduction methods such as CCA and PLS, in-sample correlations between latent factors can substantially overestimate their true generalizability^13,14^. As a result, reported molecular “signatures” may reflect model-specific artifacts rather than consistent biological patterns, particularly in studies with limited sample size and correlated features^15^. These findings highlight the importance of explicitly evaluating out-of-sample (OOS) projection performance when interpreting multivariate components.

Emerging methodological approaches only address a single facet of stability^16–19^. For example, StabilityCCA^20^ evaluates the selection stability of sparse weight vectors in CCA-based models while Stabl^21^ evaluates the selection stability of predictive biomarkers from sparse models. Neither jointly assesses the reproducibility of latent-component sample scores together with strict OOS inference. A unified framework that simultaneously evaluates stability at the level of latent-component scores and feature weights, that explicitly separates model fitting from inference, is still lacking. This gap is particularly important for multi-omics integration methods, where biological interpretation depends not only on which features are selected, but also on the reproducibility of the latent structures that link them^22^.

To address this challenge, we propose a comprehensive stability-driven framework for multi-omics integration that prioritizes reproducibility and OOS inference. Using sparse generalized canonical correlation analysis (SGCCA)^4,23,24^ (**Fig. 1a**) within a repeated cross-validation framework (**Fig. 1b**), we systematically evaluate (i) the reproducibility of latent component sample scores across resampling, (ii) the stability of feature selection and corresponding sparse weight vectors, and (iii) the consistency of associations based on strictly OOS projections (**Fig. 1c**). We further assess the feature-level stability of selected stable latent components by assessing sparse weights and feature– component correlations (loadings) (**Fig. 1d**) and perform pathway and network analyses using the resulting stable features (**Fig. 1e**). By separating model fitting from inference, this approach provides an unbiased assessment of the generalizable biological relationships captured by latent components.

**Figure 1:**
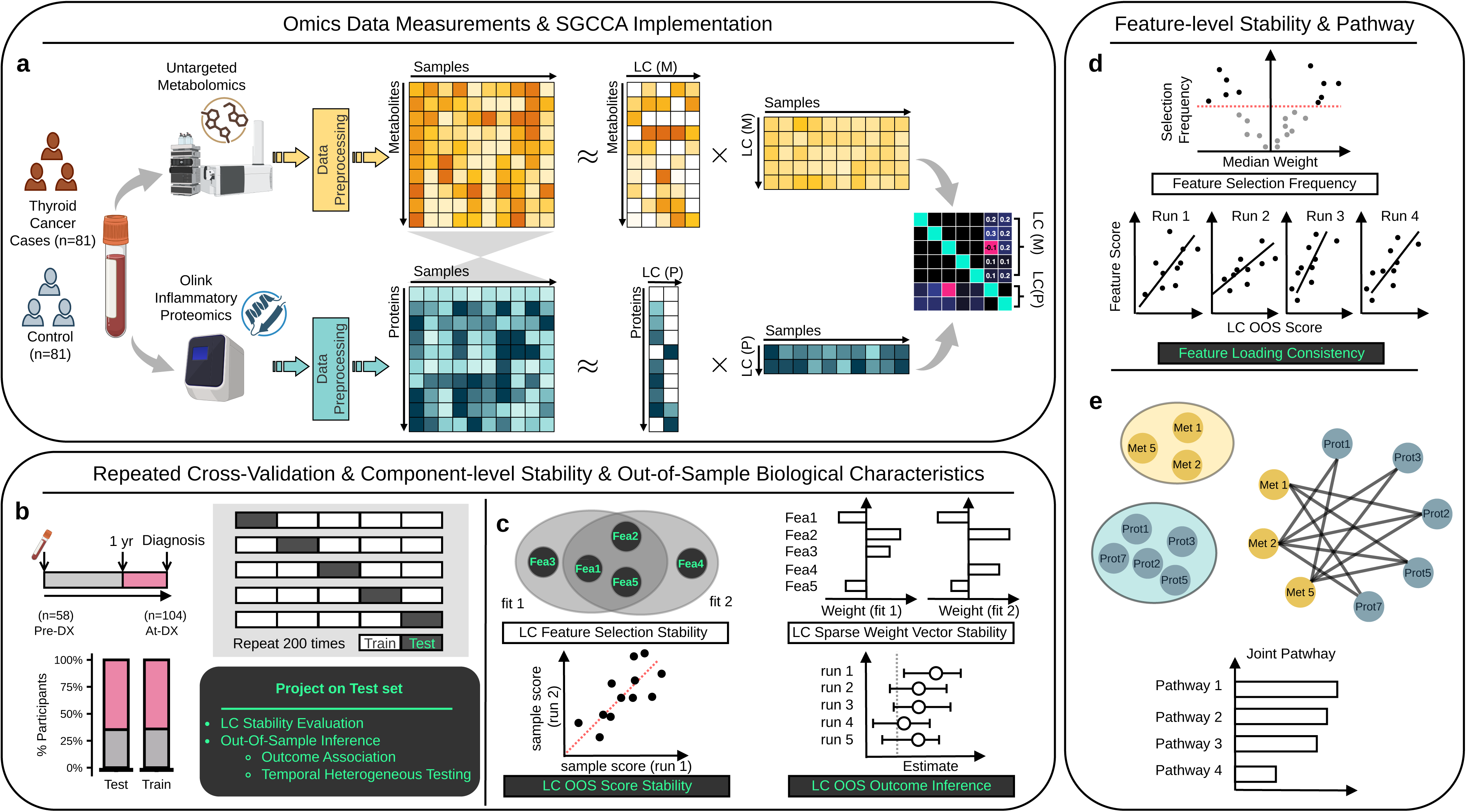
Stability-driven multi-omics integration framework overview. a. Plasma untargeted metabolomic and Olink inflammatory proteomic profiles from thyroid cancer cases (n=162 total; 81 cases, 81 controls) were integrated using sparse generalized canonical correlation analysis (SGCCA) to derive latent components (LC) capturing shared variation across omics layers. b. Within a repeated cross-validation framework (5-fold, 200 repetitions), SGCCA was fitted on training folds and projected onto held-out folds to generate strictly out-of-sample (OOS) latent component scores. c. Latent components were evaluated for stability across resampling, including reproducibility of sample scores, stability of feature selection and component sparse weight vectors, and consistency of OOS associations. d. Feature level stability was assessed using both sparse weights and loading-based correlations to identify stable molecular contributors to each component, including correlated features not retained by sparse selection. e. Stable features were used for pathway enrichment and cross-omics network analysis to support biological interpretation.

We apply this framework to integrate untargeted annotated metabolomic and Olink proteomic inflammation profiles in a thyroid cancer case–control study. Notably, our goal is not to develop a predictive model or identify individual biomarkers, but rather to evaluate the reliability of multi-omics integration results under realistic sample size constraints. Previous analyses of this cohort have reported associations between environmental exposures^25^, inflammation proteins^26^, and disease risk. However, such single-omics approaches are limited in their ability to capture shared molecular variation across biological layers. By integrating multiple omics layers within a stability-driven framework, we identify reproducible latent structures that reflect cross-omics biological processes while avoiding overinterpretation of unstable components.

## Results

SGCCA was applied jointly on metabolomics and inflammatory protein data to extract latent components capturing coordinated cross-omics variation (**Supplementary Method 1**). All model-related stability evaluation results are based on 1,000 model fits (5-fold, 200 repetitions); all OOS related evaluation results are based on stacked 5-fold scores, 200 repetitions. Model hyperparameters (the number of latent components per block and the per-component sparsity) were selected by the repeated cross-validation tuning procedure (**Supplementary Method 1**) and remained fixed across all resampling iterations to ensure comparability of components. This procedure selected eight latent components for metabolomics (LC1 [M]–LC8 [M]) and three for targeted proteomics (LC1 [P]–LC3 [P]), with sparsity parameters retaining approximately 45% of metabolites and 60% of inflammation proteins per component (**Supplementary Fig. 1**). As an example of a single model fit using the selected hyperparameters, SGCCA revealed structured cross-omics correlation patterns between metabolomics and proteomics blocks (**Supplementary Fig. 2**). We next evaluated the reproducibility of these latent components across repeated cross-validation.

### Stability-driven evaluation identifies reproducible latent components

We first evaluated the reproducibility of latent components derived from SGCCA across repeated cross-validations (**Fig. 2**, **Supplementary Table 1**). Visualization of latent component (LC) scores across subjects revealed consistent structures for the lower-order component in both metabolomics and proteomics, whereas higher-order components showed inconsistent patterns (**Fig. 2a**). Pairwise correlations of component scores across resampling of metabolomics LCs showed that LC1 had the most stable signal (**Fig. 2b, f**) with a median Spearman’s ρ = 0.59 (IQR = [0.58, 0.60]). LC2 (M) was weaker (median = 0.47, IQR = [0.45, 0.49]) followed by LC3–LC8 (median = [0.24, 0.09]). Similarly for inflammation proteins, LC1 (P) exhibited high reproducibility with a median Spearman’s ρ = 0.76 (IQR = [0.75, 0.77]) across resampling iterations, while subsequent components showed lower stability (LC2: Spearman’s ρ = 0.60, IQR = [0.67, 0.72]; LC3: Spearman’s ρ = 0.50, IQR = [0.47, 0.54]). Sparse weight profiles indicated that only a subset of features were consistently selected across resampling iterations (**Fig. 2c, g**). Selected metabolite features spanned multiple chemical superclasses, with lipids and lipid-like molecules and organic acids and derivatives representing the largest contributing classes as illustrated by the overall metabolomic feature space composition (**Fig. 2d**). Lipids and lipid-like molecules and organic acids and derivatives represent the largest chemical classes. Among all extracted components, LC1 (M), LC1 (P) and LC2 (P) demonstrated the most stable selection patterns and directional consistency across runs (sparse weight vector Spearman’s p LC1 (M) median = 0.67, IQR = [0.64, 0.71]; LC1 (P) median = 0.91, IQR = [0.88, 0.93] and LC 2(P) median = 0.76, IQR = [0.73, 0.79]). In contrast, higher-order components displayed substantial variability in selected features, reflecting sensitivity to sampling variability. Additionally, feature selection stability summarized by the Nogueira index was highest for LC1 (M) (0.4) and declined for subsequent metabolomic components (0.27–0.03) (**Fig. 2h**). Similarly, LC2 (P) had the highest Nogueira index (0.52), followed by LC1 (P) (0.37) and LC3 (P) (0.122). LC1 (M), LC 1(P), and LC 2(P) achieved both high subject-level reproducibility and feature-level stability in joint visualization of score reproducibility and weight stability (**Fig. 2i**). Therefore, only a limited number of the latent components, namely LC1 (M), LC1 (P), and LC2 (P), captured reproducible cross-omics variation, whereas most components were unstable and likely reflected sampling-dependent variability. This highlights the importance of stability-based filtering prior to biological interpretation.

**Figure 2:**
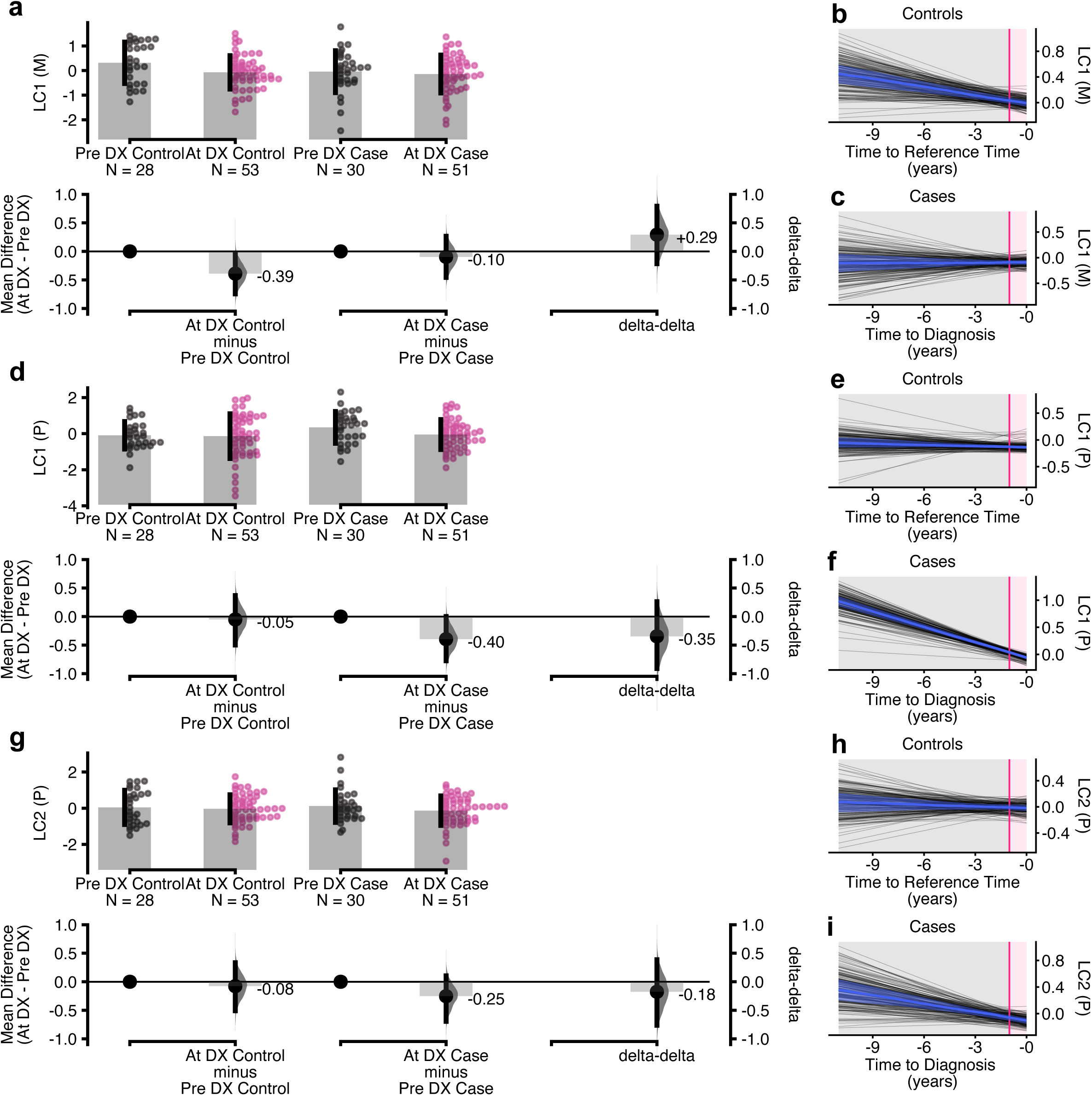
Stability evaluation distinguishes reproducible from unstable latent components. a. Heatmaps of latent component (LC) scores across subjects (n=162) for metabolomics (LC1–LC8 (M)) and inflammatory proteomics (LC1–LC3 (P)). b. Distribution of pairwise Spearman correlations of latent component scores across resampling iterations, summarizing subject–level score reproducibility for each component. c. Heatmap of metabolite selection and corresponding sparse weights across resampling for each latent component (LC1–LC8 [M]). Blocks represent components, rows represent resampling iterations, and columns represent metabolites. Side panels summarize the superclass distribution of features with positive (right) and negative (left) weights for each component, with colors corresponding to metabolite superclasses. d. Pie chart illustrating the chemical composition of the 284 metabolomic feature space by superclass. e. Heatmap of proteomic loadings across latent components (LC1–LC3 [P]), with corresponding summaries of selected features with positive and negative weights. f. Stability of latent component scores across resampling, measured by Spearman correlation, summarized per component. g. Stability of sparse weight vectors across resampling, reflecting variability in feature selection and contribution across components. h. Summary of feature selection stability for each component, including selection rate (median, minimum, and maximum) and Nogueira index. i. Joint visualization of score reproducibility (x-axis) and weight stability (y-axis) for each component, illustrating the relationship between subject-level stability and feature-level stability.

### Distinct disease-associated temporal patterns in reproducible out-of-sample latent components

We next assessed whether the latent components exhibited reproducible associations with disease status using strict OOS projections across repeated cross-validation runs (**Supplementary Table 2**). This was tested in the subgroups of pre-diagnosis (sampled 1–8 years before diagnosis and their matched controls), at-diagnosis (sampled < 1 year before diagnosis and their matched controls), as well as the combined analysis. Overall, LC1 (P) demonstrated the most consistent positive association with disease status, particularly in the pre-diagnostic subgroup, with a median OOS odds ratio (OR) = 1.85 (IQR = [1.67, 1.98]) and 99.0% of cross-validation runs with positive effect estimates. A positive directional for LC1 (P) was also observed at-diagnosis (86.0% positive) and in the combined analysis (96.0% positive), indicating stability of the association across temporal strata and resampling iterations. Resampling with subject-level averaged OOS scores for LC1 (P) were consistent with this. The largest case-control difference for LC1 (P) was observed in the pre-diagnostic subgroup compared to the at-diagnosis subgroup or the combined analysis (**Supplementary Fig. 3**), supporting the presence of a reproducible early disease-associated inflammation proteomic signal. LC2 (P) showed weaker and less stable associations overall, with mixed directions across subgroups and comparatively modest separation between cases and controls. The metabolomic component LC1 (M) exhibited negative associations across resampling iterations, particularly in the pre-diagnostic subgroup (median OR = 0.73, IQR = [0.65, 0.82], 96.5% negative) and in the combined analysis (92.5% negative). Correspondingly, averaged OOS scores demonstrated lower LC1 (M) values among cases relative to controls within the pre-diagnostic subgroup (**Supplementary Fig. 3**), supporting a reproducible negative association with disease status. Together, these analyses support the robustness and reproducibility of the disease-associated patterns of LC1 (P) and LC1 (M). Similar patterns were observed in sensitivity analyses restricted to complete matched case-control pairs (n = 152) (**Supplementary Fig. 4**).

Having established reproducible disease-associated signals, we next examined whether these components exhibited distinct temporal dynamics between samples collected pre-diagnostic and at-diagnosis periods and their matched controls (**Fig. 3**). LC1 (M) exhibited a distinct temporal pattern among controls. Controls showed lower LC1 (M) scores at diagnosis/reference relative to the pre-diagnostic period (mean difference = - 0.38, BCa 95% CI = [–0.75, –0.02], permutation p = 0.030; **Fig. 3a**, **Supplementary Table 3**), together with a consistent negative temporal relationship over time (median slope = –0.04, IQR = [–0.06, –0.03]; **Fig. 3b**, **Supplementary Table 3**), indicating a baseline temporal metabolomic trajectory. In comparison, disruption or attenuation of this underlying temporal metabolic pattern was observed in individuals who developed thyroid cancer; LC1 (M) trajectories among cases were comparatively weak and less consistent (median slope = 0.01, IQR = [–0.02, 0.03]; **Fig. 3c**, **Supplementary Table 3**). A contrasting trend was observed in LC1 (P), where lower LC1 (P) scores were observed at-diagnosis relative to the pre-diagnostic period (mean difference = –0.40, BCa 95% CI = –0.78, –0.003], permutation p = 0.045; **Fig. 3d**, **Supplementary Table 3**). Trajectory analyses demonstrated a robust negative relationship between LC1 (P) scores and time to diagnosis among cases (median slope = –0.10, IQR = [–0.11, –0.09]; **Fig. 3f**, **Supplementary Table 3**), whereas controls exhibited comparatively minimal temporal change (median slope = –0.01, IQR = –0.02, 0.001]; **Fig. 3e**, **Supplementary Table 3**). These findings suggest that LC1 (P) captured a progressive proteomic trajectory associated with thyroid cancer development. In LC2 (P), cases showed moderate negative temporal trajectories (median slope = –0.05, IQR = [–0.06, –0.03]; **Fig. 3i**), but estimation plots revealed no clear separation between pre-diagnostic and at-diagnosis samples (**Fig. 3g**), consistent with the weaker disease-association signals observed in the OOS analyses.

**Figure 3:**
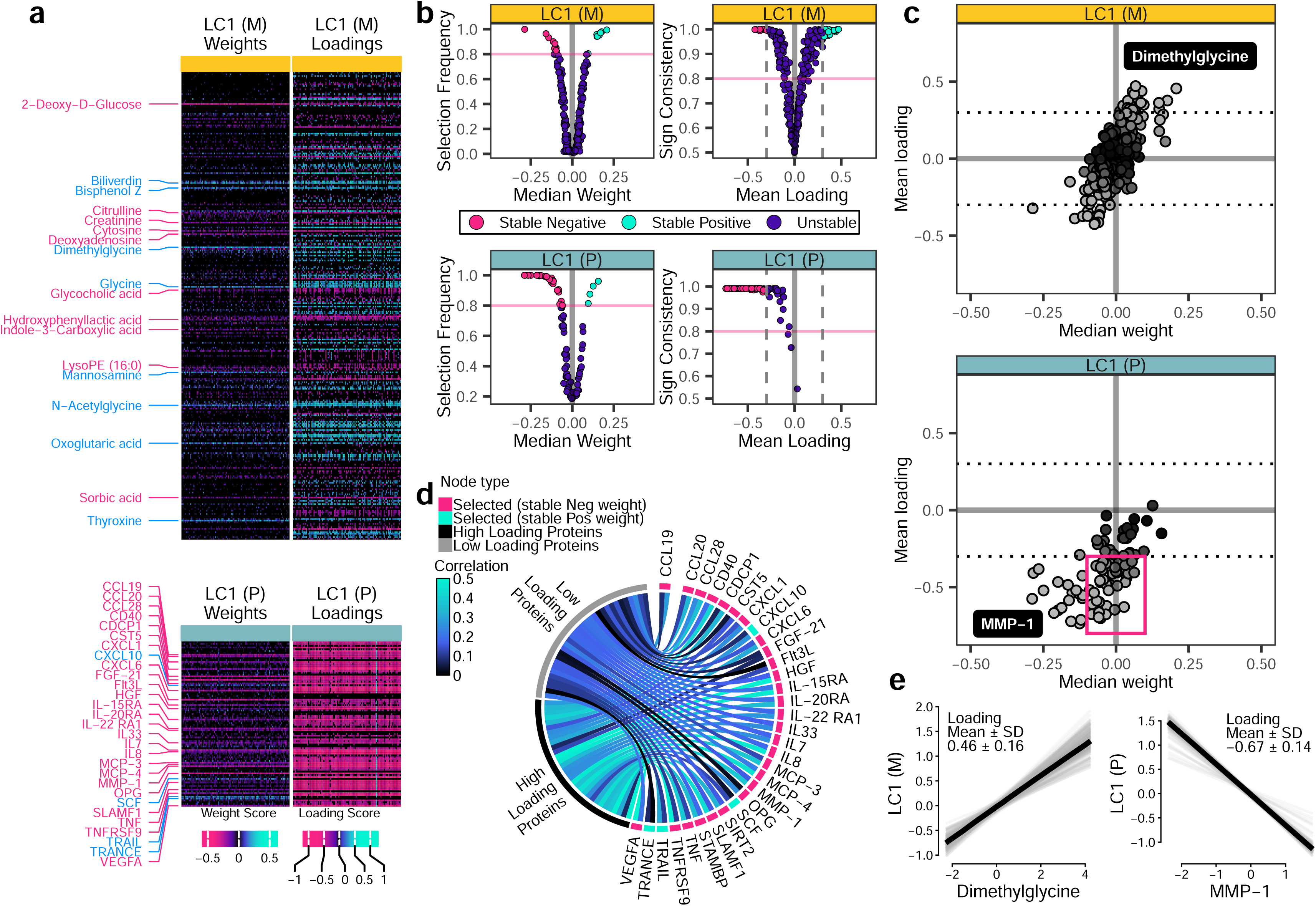
Out-of-sample biological characteristics of stable latent components. a, d, g. Estimation plots of differences in LC scores between pre-diagnosis and at-diagnosis samples for LC1 (M) (a), LC1 (P) (d), and LC2 (P) (g). Points represent individual samples, and mean differences (at-diagnosis – pre-diagnosis) are shown with bias-corrected and accelerated 95% bootstrap confidence intervals (5,000 resamples). The delta-delta contrast represents the difference in temporal change between cases and controls (case [at-diagnosis - pre-diagnosis] – control [at-diagnosis - pre- diagnosis]). Group sizes: pre-diagnostic controls n = 28, at-diagnostic controls n = 53, pre-diagnostic cases n = 30, at-diagnostic cases n = 51. b, c, e, f, h, i. Temporal analysis of LC scores as a function of time to diagnosis (cases) or matched reference time (controls). Panels are organized by components: LC1 (M) (b, c), LC1 (P) (e, f), and LC2 (P) (h, i), with matched controls (b, e, h) preceding cases (c, f, i). Thin lines represent linear fits from individual cross-validation runs, illustrating model variability, while the solid line indicates the median fitted trajectory, and shaded areas represent the interquartile range (25th–75th percentiles) across runs. Time is expressed in years before diagnosis (cases) or matched reference time (controls), with values closer to zero meaning closer to the diagnosis/reference point.

### Feature-level stability reveals coordinated molecular structure within selected LCs

We next evaluated the stability of individual feature contributions within the stable LC1 (M) and LC 1 (P) components across repeated cross-validation runs (**Fig. 4**). SGCCA weights quantify conditional feature contributions used to construct the latent component, while component loadings revealed a broader set of features showing stable correlations with the derived components. (**Fig. 4a**, **4b**). Sparse weight profiles identified 19 metabolites and 31 proteins as consistently selected features for LC1(M) and LC1(P) respectively. Beyond the stably selected features, loading-based analyses showed that a broader set of features, particularly for LC1 (P), exhibited consistent directions and magnitudes of association with the latent components (**Fig. 4c**). Loading-based analyses additionally identified 28 metabolites and 39 proteins with stable loadings exceeding ±0.3 but not directly selected by the sparsity penalty. For LC1 (M), 19 selected features showed mean absolute loadings of 0.33–0.46 while additional 28 non-selected features but with stable loadings exhibited comparable loading magnitudes 0.3–0.47 (**Supplementary Table 4**). Notably, for LC1 (P), 39 features with stable loadings but not consistently selected by the model (high loading features, **Fig. 4d**, **Supplementary Fig. 6**) showed stronger correlations with the 31 selected features (median Spearman’s ρ = 0.28, IQR = [0.18, 0.39]) than remaining 22 features with unstable loadings (low loading features, **Fig. 4d**, **Supplementary Fig. 6**; median Spearman’s ρ = 0.14, IQR = [0.07, 0.23]).

**Figure 4:**
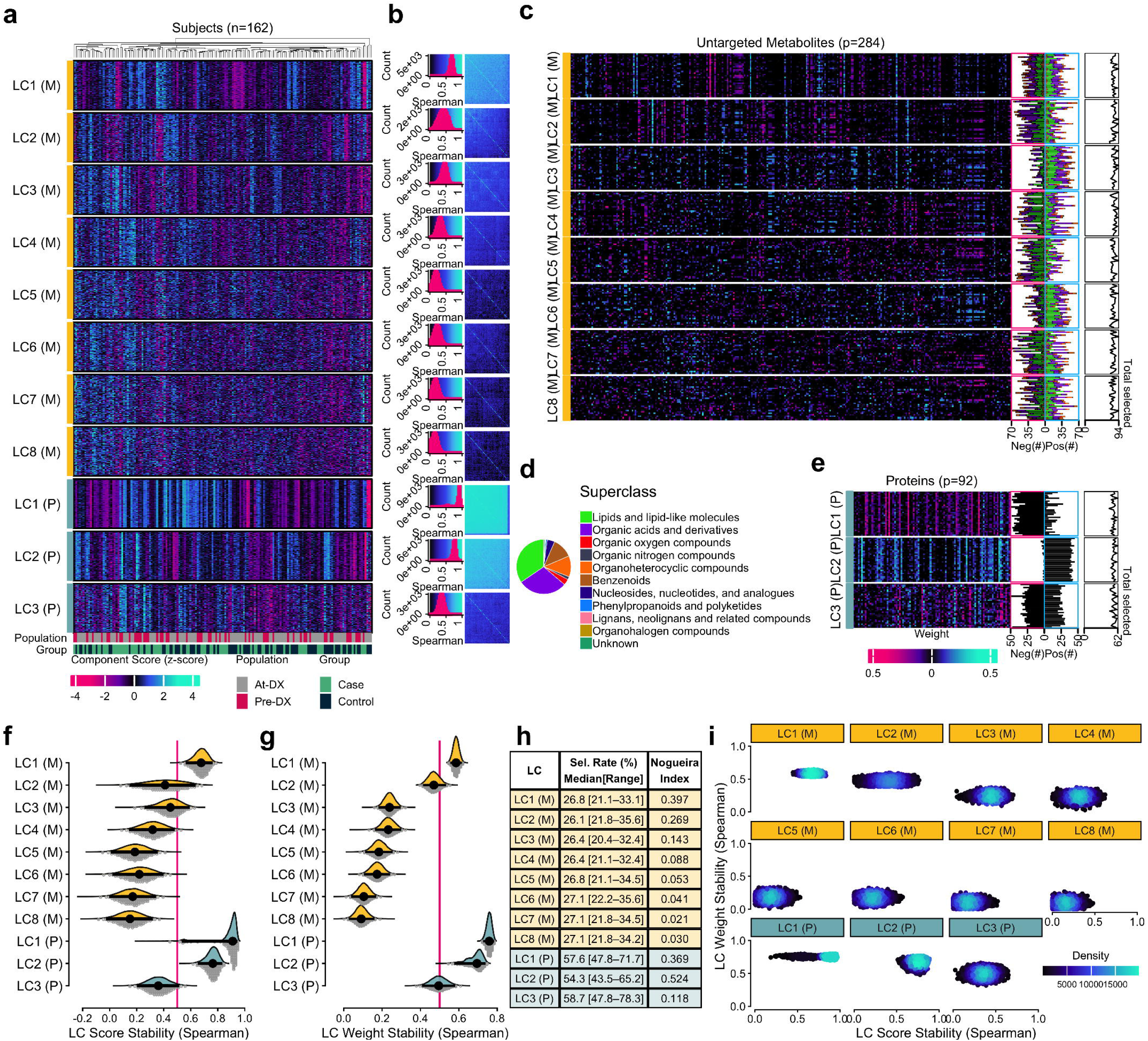
Feature-level stability of sparse weights and loadings for LC1 (M) and LC1 (P) a. Heatmaps of feature weights (left) and feature - component correlations (loadings; right) across resampling iterations for LC1 (M) (top) and LC1 (P) (bottom). Rows represent individual features and columns represent resampling iterations. Color indicates the magnitude and direction of weights or loadings. Row labels on the left side highlight the 19 and 31 features with stable sparse weights for LC1(M) and LC1(P), respectively. b. Feature selection frequency and sign consistency across resampling iterations for LC1(M) (top) and LC1(P) (bottom). Selection frequency is defined as the proportion of resampling iterations in which each feature was assigned a non-zero sparse weight (x-axis). Sign consistency reflects the directional agreement of non-zero weights across iterations (y-axis). Vertical and horizontal reference lines at 0.8 indicate thresholds used to identify features with stable selection and directional consistency. Vertical reference lines at ±0.3 on the loading axis indicate the threshold for stable feature - component associations. c. Scatter plot of mean loading values (y-axis) versus median sparse weights (x-axis) across resampling iterations for LC1 (M) (top) and LC1 (P) (bottom). Each point represents one feature. Horizontal dotted reference lines at ±0.3 indicate the loading magnitude threshold used to identify features with stable feature-component associations. Labeled features represent examples with the largest concordant weight and loading magnitudes, Dimethylglycine for LC1(M) and MMP-1 for LC1(P). The rectangle in LC1(P) highlights representative proteins with low sparse weights but stable high loadings, which are incorporated into the chord diagram in panel d as high loading features. d. Chord diagram illustrating the correlation structure between proteins. Node colors represent consistently selected features with stable positive weights (cyan), consistently selected features with stable negative weights (magenta), non-selected features with high stable loadings (black), and remaining features (grey). Link colors represent the direction and magnitude of Spearman correlations between features. e. Stable feature - component linear relationships for representative selected features contributing to LC1(M) (Dimethylglycine) and LC1(P) (MMP-1). Each line represents the fitted linear relationships between z-scaled feature abundance (x-axis) and z-scaled component scores (y-axis) estimated across resampling iterations. The black line indicates the median fitted relationship, and colored lines correspond to individual resampling iterations, with color reflecting the direction and strength of the Spearman correlation between the feature and the component across iterations.

Together, these findings indicate that sparse feature selection captured a minimal but reproducible feature set, while loading-based analyses revealed a broader coordinated molecular structure associated with the latent components. This distinction reflects a fundamental property of sparse integration methods: because SGCCA enforces conditional feature selection, correlated features may be systematically excluded despite exhibiting reproducible alignment with the latent molecular axis. Detailed feature-level stability metrics including selection frequency, weight distributions, and loading consistency for LC1(M) and LC1(P) are provided in **Supplementary Tables 4 and 5**, and corresponding results across all remaining latent components are summarized in **Supplementary Figure 5**.

### Integrating weights and loadings in multi-omics biological interpretation

To inform the molecular structure of all stable components, we compared sparse feature selection with loading-based feature characterization across LC1 (M) and LC1 (P) (**Fig. 5**). First, we highlight differences in the number of metabolite-inflammation protein associations when restricted to the 50 features selected by sparse weights only (**Fig. 5a**) compared to the denser network of correlated metabolite-inflammation protein interactions when incorporating an additional 67 features with stable loadings (**Fig. 5b**). Forest plots of case-control odds ratios confirmed that loading-selected features exhibited disease associations consistent in direction and magnitude with weight-selected features, supporting their biological relevance to thyroid cancer risk (**Fig. 5a, b**). A metabolite-inflammation protein correlation network using pairwise Spearman correlations (FDR-adjusted p<0.05) identified several high-centrality metabolites and inflammation proteins that may function as bridge nodes connecting metabolic and inflammation pathways (**Supplementary Fig. 7**). Notably, glutarylcarnitine and 5-hydroxylysine, identified through stable loading-based selection rather than sparse weights alone, exhibited high network centrality, suggesting that these metabolites may function as key intermediates linking inflammation and metabolic alterations in thyroid cancer (**Supplementary Fig. 7**). Most metabolites exhibited correlations of consistent direction with inflammation proteins-either predominantly positive or predominantly negative-suggesting that individual metabolites tend to be coordinately associated with the inflammation proteome in a uniform direction. In contrast, individual inflammatory proteins showed more heterogeneous correlation patterns across metabolites, with both positive and negative associations present simultaneously, reflecting the broader and more diverse functional roles of inflammatory proteins across metabolic pathways (**Supplementary Fig. 7**). Joint pathway over-representation analysis using IMPaLA revealed that inclusion of loading-selected features substantially expanded pathway coverage compared to sparse weight-selected features alone. Using stable weights only, nine proteins overlap with GPCR-related pathways, with no significant joint enrichment observed (joint p-values = [0.13, 0.88]). In contrast, the expanded feature set combining stable weights and stable loadings yielded significant joint enrichment across the GPCR signaling hierarchy, including Signaling by GPCR (joint p = 0.009), GPCR ligand binding (joint p = 0.006), GPCR downstream signaling (joint p = 0.020), Class A/1 Rhodopsin-like receptors (joint p = 0.003), and G alpha (q) signaling events (joint p = 0.025), with overlapping inflammation proteins increasing from 9–10 to 15–19 and overlapping metabolites increasing from 0–1 to 8–9 per pathway (**Fig. 5c**). Several enriched Reactome pathways representing hierarchical GPCR-related signaling processes with highly overlapping molecular members were interpreted collectively as a coordinated GPCR signaling module (**Fig. 5d, Supplementary Table 6**). These results suggest that loading-based feature expansion recovers coordinated multi-omics pathway structure that would otherwise be missed by sparse weight selection alone, particularly for GPCR-mediated inflammatory signaling pathways relevant to thyroid cancer biology.

**Figure 5:**
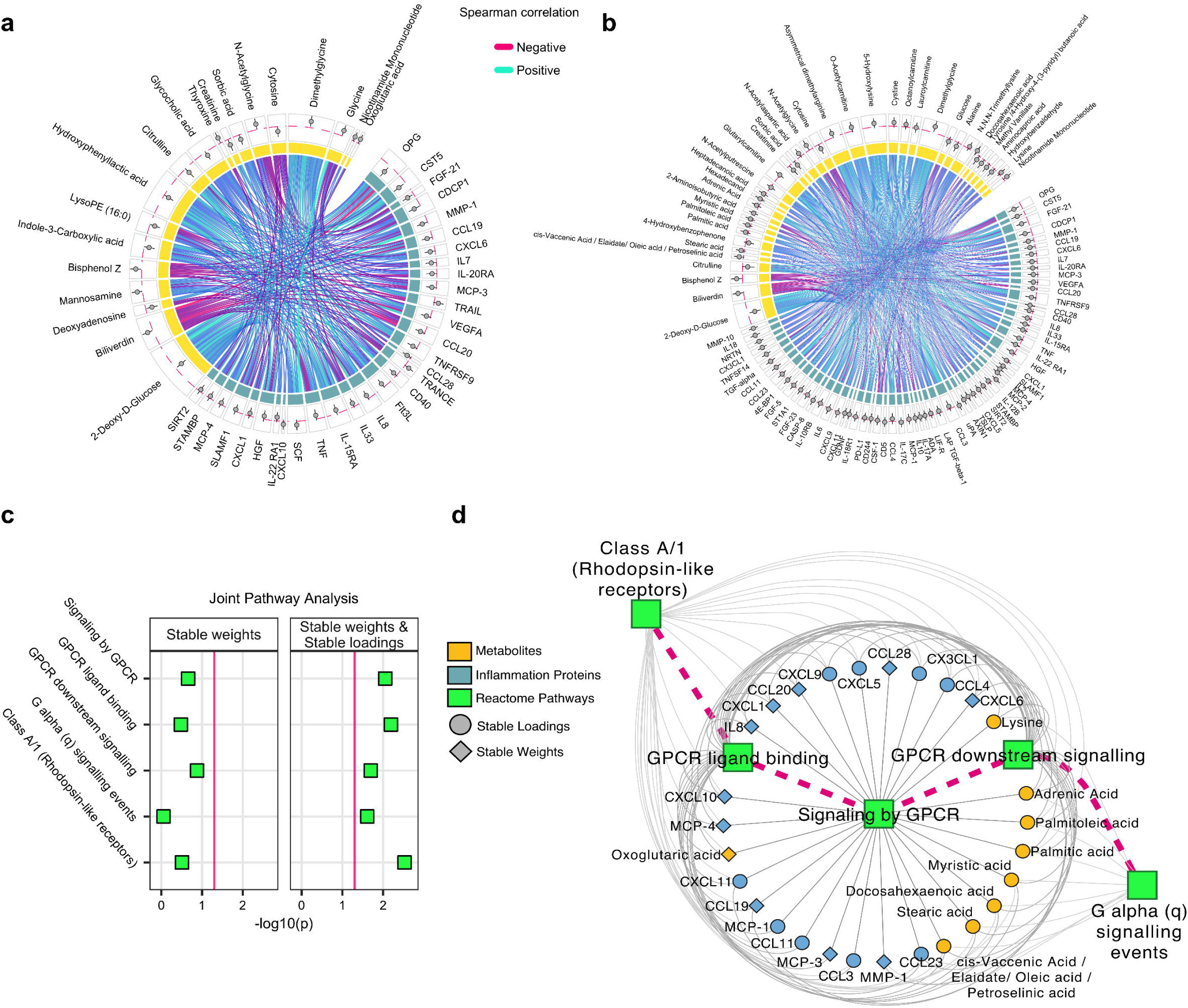
Cross-omics network and joint pathway enrichment for stable LC1 (M) and LC1 (P) features. a. Circos plot illustrating pairwise Spearman correlations between metabolite and protein features selected based on stable sparse weights in LC1(M) and LC1(P). Links represent pairwise correlations, with line width proportional to absolute correlation strength and color indicating direction (blue: positive, magenta: negative). Forest plots display odds ratios (ORs) with 95% confidence intervals from logistic regression models adjusting for age, sex, BMI, race/ethnicity, and sample storage time for each selected feature. b. Circos plot illustrating pairwise Spearman correlations among the expanded feature set comprising both stable weight-selected features and additionally identified stable loading-selected features for LC1(M) and LC1(P). Links represent pairwise correlations, with line width proportional to absolute correlation strength and color indicating direction (blue: positive, magenta: negative). Forest plots display odds ratios (ORs) with 95% confidence intervals from logistic regression models adjusting for age, sex, BMI, race/ethnicity, and sample storage time for each selected feature. c. Joint pathway enrichment analysis results based on LC1 features selected using stable weights only and on the combined feature set with stable weights and consistently high loadings. Each point represents a pathway, with size proportional to the number of matched features and color indicating statistical significance (-log10 joint enrichment p-value). A magenta solid vertical reference line indicates the nominal significance threshold (joint p = 0.05). d. Feature-pathway network illustrating the recovery of coordinated GPCR-related multi-omics pathway structure following inclusion of loading-associated features. Inflammation proteins and metabolites are connected to their overlapping IMPaLA pathways, with Reactome pathway hierarchy relationships incorporated to represent GPCR signaling pathway structure. Node color indicates molecular type (yellow: metabolites, blue: inflammatory proteins, green: pathways) and node shape indicates feature-selection category (diamonds: stable weight-selected features, circles: stable loading-selected features). Grey edges connect individual metabolite and protein features to their corresponding pathways, indicating feature-to-pathway membership; magenta edges represent hierarchical relationships between Reactome pathways within the GPCR signaling module.

## Discussion

### Stability-driven multi-omics integration identifies reproducible disease-associated latent structures

In this study, we present a stability-driven framework for multi-omics integration that prioritizes reproducibility and strict OOS evaluation. Using repeated cross-validation and resampling-based stability assessment, we demonstrate that only a subset of latent components derived from sparse multiblock models represent reproducible biological structure, whereas higher-order components are often unstable and sensitive to sampling variability. Rather than emphasizing associations from a single fitted model, our framework focuses on latent molecular structures that remain consistent across repeated resampling and OOS projection. Importantly, the objective of this framework is biological reproducibility rather than predictive optimization, with the goal of identifying molecular structures that generalize beyond a single model fit. Applying this framework to a study of thyroid cancer, we identified a reproducible latent axis linking inflammation proteomic and metabolomic variation. The inflammation proteomic component LC1 (P) demonstrated the strongest and most reproducible disease-associated signal, particularly during the pre-diagnostic period, suggesting progressive inflammation remodeling prior to clinical diagnosis. In contrast, the metabolomic component LC1 (M) exhibited temporal patterns primarily among controls, with attenuation of these trajectories among individuals who subsequently developed thyroid cancer. Together, these findings suggest that disease-associated molecular variation may be dominated by a limited number of coordinated inflammation and metabolic structures.

### Reproducibility and out-of-sample inference as prerequisites for biological interpretation

A central finding of this study is that reproducibility should be considered a prerequisite for biological interpretation in multi-omics integration. Higher-order latent components frequently failed to replicate across resampling iterations, consistent with theoretical expectations in high-dimensional settings where correlated features, sparse optimization^12,15^, and moderate sample size contribute to non-identifiability and instability of latent structure. Importantly, instability should not be viewed solely as a methodological limitation, but rather as informative evidence regarding the reliability of inferred biological signals. In this context, OOS inference provides a more reliable basis for interpretation than in-sample statistical significance alone. While individual resampling iterations yielded variable effect estimates, consistent directional effects across repeated OOS evaluations supported the reproducibility of stable latent components. These findings align with broader statistical perspectives emphasizing that single-model p-values may be unstable in high-dimensional settings, whereas repeated resampling provides a more robust assessment of generalizability and sampling variability. OOS reproducibility is particularly important in multi-omics studies because latent components may otherwise reflect cohort-specific correlation structure rather than persistent biological structures.

### Integrating sparse weights and loading stability improves biological interpretation

Our results further demonstrate that feature-level stability is essential for meaningful biological interpretation of latent molecular structure. Sparse SGCCA weights identified a minimal subset of conditionally selected features contributing directly to component construction, whereas loading-based analyses captured a broader set of correlated features that aligned with the same latent molecular axis. Because sparse optimization may arbitrarily retain one feature among correlated groups, biologically relevant features can remain reproducibly associated with the latent component despite inconsistent sparse selection. Joint evaluation of sparse weight stability and loading consistency, therefore, captured both the minimal predictive structure and broader, coordinated molecular structures. This distinction proved particularly important for downstream pathway and network analyses, where incorporating stable loading-based features revealed substantially richer cross-omics relationships and biological pathways than sparse feature selection alone.

Sparse feature selection identifies the minimal discriminative molecular signature, whereas inclusion of high-loading latent-component features improves systems-level biological interpretability by recovering coordinated pathway structure across omics layers. Feature inclusion was based on reproducibility criteria derived from resampling stability rather than nominal statistical significance. Inclusion of high-loading latent-component features improved recovery of coordinated pathway-level molecular structure, resulting in stronger multi-omics convergence within biologically relevant GPCR signaling pathways.

### Broader implications for high-dimensional multi-omics studies

More broadly, these findings suggest that instability is an inherent characteristic of high-dimensional multi-omics that needs to be quantified and accounted for within the analytical workflow. By treating reproducibility and OOS validity as primary criteria for interpretation, the proposed framework provides a generalizable strategy for improving the reliability of multi-omics inference while reducing the risk of model-dependent biological conclusions. Importantly, our findings do not suggest that multi-omics integration is ineffective in moderate-sized studies. Rather, biologically reproducible signals in heterogeneous human populations may exhibit modest effect sizes and substantial uncertainty. In this setting, inferred latent structure must remain reproducible across repeated resampling and generalizable beyond a single fitted model.

### Limitations and future directions

Several limitations should be considered. First, the moderate sample size relative to the dimensionality of the omics data remains an inherent challenge. However, this setting reflects common conditions in biomedical multi-omics studies, where high-dimensional measurements are frequently collected from limited cohorts. Our findings, therefore, emphasized the importance of explicitly evaluating reproducibility under realistic sample size constraints. Next, we applied a mean absolute loading cutoff of 0.3 across resampling iterations, chosen to reflect a moderate and interpretable feature-component association. However, no consensus criterion currently exists for this threshold in multi-omics integration, and alternative approaches such as rank-based or proportion-based may yield different feature sets and downstream pathway findings. Another limitation of the current stability assessment is that the number of latent components per block and the sparsity hyperparameters were determined in a prior repeated cross-validation tuning step and subsequently held constant throughout the stability evaluation. Fixing the model specification was necessary to ensure that component-, weight-, and feature-level stability could be assessed consistently across iterations, as models fitted with different numbers of components or sparsity settings would produce non-comparable latent structures. Consequently, the reported stability estimates reflect variability conditional on the selected model specification and do not capture the additional uncertainty that may arise from re-tuning hyperparameters within each resampling iteration. Evaluating stability under a fully nested tuning framework, where hyperparameters are re-optimized at every resampling step, represents an important avenue for future methodological research. In addition, the study was conducted within a single cohort, and external validation in independent populations will be necessary to confirm generalizability^27^. Finally, this study focused on SGCCA as a representative multiblock integration framework while alternative integration approaches may differ in their stability properties and biological sensitivity. Therefore, future work will extend this framework across three complementary dimensions of reproducibility: population reproducibility through validation in independent cohorts, algorithmic reproducibility through application to alternative integration methods, and biological reproducibility through incorporation of additional omics modalities. These extensions will determine whether identified latent molecular programs remain stable across datasets, analytical frameworks, and biological contexts.

### Conclusion

In summary, we present a stability-driven multi-omics integration framework that combines latent component reproducibility, feature-level stability assessment, and strict OOS inference to identify robust latent molecular structure. Our findings demonstrate that only a subset of latent components exhibit reproducible biological structures and generalizable disease-associated patterns, whereas higher-order components are often unstable and prone to overinterpretation. By prioritizing reproducibility, uncertainty quantification, and OOS validity, this framework provides a practical strategy to improve reliability and interpretability of high-dimensional multi-omics analyses.

## Methods

### Study population

The study cohort was drawn from the BioMe Biobank, a medical record–linked repository at the Icahn School of Medicine at Mount Sinai. BioMe has enrolled over 60,000 participants since 2007, representing a racially and socioeconomically diverse population in New York City. The cohort for this study included 88 thyroid cancer cases with plasma collected at or prior to diagnosis and 88 matched controls without cancer. Controls were matched on age (±5 years), sex, race/ethnicity, body mass index (BMI), smoking status, and year of sample collection. Cases were identified by ICD-9 (193) and ICD-10 (C73) codes. Detailed cohort design, biospecimen collection, and matching procedures have been described previously^25,26^. Of the 176 matched participants (88 cases, 88 controls), 162 had complete metabolomic and Olink inflammation proteomic data after quality control; of these, 152 remained as complete matched case–control pairs and were used for the paired sensitivity analysis (**Supplementary Method 2**). Baseline characteristics of the study population are summarized in **Supplementary Table 7**. Briefly, no significant differences were observed in age, sex, BMI, or race/ethnicity between pre-diagnosis (defined as participants with plasma samples collected 1–8 years before diagnosis and matched controls) and at-diagnosis (defined as participants with plasma samples collected < 1 year before diagnosis and matched controls) populations. The cohort was predominantly female (83%), with a mean (SD) age of 46 (15.3) years at blood collection. Plasma samples were collected between 2008 and 2021, with time to diagnosis among cases ranging from 0 to 8.5 years.

### Omics data acquisition and preprocessing

Olink inflammation proteomic and metabolomic data acquisition and preprocessing procedures are detailed in prior publication^26^ and **Supplementary Method 2**. Briefly, inflammation proteins were quantified using the Olink Target 96 Inflammation panel and untargeted metabolomics profiling used liquid chromatography–high resolution mass spectrometry (LC–HRMS) in complementary chromatographic modes. We applied standard quality control, normalization, and filtering procedures within each omics block.

### Repeated cross-validation SGCCA framework

We applied sparse generalized canonical correlation analysis (SGCCA) to integrate metabolomic and proteomic data and derive latent components capturing shared cross-omics variation. The primary objective of this framework was not predictive optimization, but identification of reproducible latent molecular structure that remained stable across repeated out-of-sample resampling.

We performed model tuning and estimation within a repeated cross-validation framework (5-fold, 200 repetitions; full details are provided in **Supplementary Method 1**). An initial tuning step fixed the number of components per block to enable comparability across resampling iterations. We performed all preprocessing steps exclusively within training folds to prevent information leakage. Specifically, in the training data, we used linear models to estimate the residuals regressing age, sex, BMI, race/ethnicity, and sample storage time on each metabolomic and proteomic feature.

We then standardized features using training-derived parameters and applied both transformations to held-out samples using the corresponding training estimates. For each resampling iteration, we fitted SGCCA models in each of the five training folds. Held-out samples were then projected onto the fitted latent space to obtain their latent component scores. We stacked out-of-sample (OOS) component scores from held-out samples across folds within each repetition, ensuring complete separation between model fitting and downstream inference. Model-derived quantities (e.g., sparse weights and feature selection) were evaluated within training folds, whereas all downstream association analyses and temporal trajectory analyses used the stacked OOS component scores.

### Stability evaluation of latent components

Latent components were evaluated along three complementary dimensions. First, score stability, reflecting reproducibility of the latent representation, was assessed by examining the consistency of stacked OOS component scores across repeated resampling iterations using Spearman correlation. Second, selection stability, reflecting reproducibility of sparse feature selection, was quantified using the Nogueira stability index^28^, which measures agreement in selected features across model fits. Third, weight stability, reflecting preservation of relative feature contribution within the latent component, was evaluated by assessing similarity of sparse weight vectors across resampling iterations using correlation-based metrics (see **Supplementary Table 8**).

### Out-of-sample association and temporal analyses

We evaluated associations between latent component scores and thyroid cancer status using regression models fitted exclusively on stacked OOS component scores. Case-control analyses used logistic regression, whereas temporal analyses used linear regression. We computed effect estimates separately within each cross-validation repetition and summarized using robust statistics, including the median, interquartile range (IQR), and directional consistency across runs. We evaluated temporal dynamics of latent component scores relative to time to diagnosis (cases) or matched reference time (controls) using linear regression models fitted on OOS component scores within each resampling iteration. Temporal effect estimates were summarized across repetitions as median slopes, IQRs, and directional consistency.

Group differences in latent component scores were visualized using estimation statistics implemented in the dabestr package (version 2025.3.15), following estimation-focused approaches recommended by Ho et al^29^. Bootstrap confidence intervals for mean differences came from nonparametric subject-level bootstrap resampling (5,000 resamples) with bias-corrected and accelerated (BCa) intervals.

As a sensitivity analysis, estimate comparisons were additionally repeated within the subset of complete matched case-control pairs remaining after quality control (n = 152). Paired differences in OOS latent component scores were evaluated using subject-level paired estimation statistics with 5,000 nonparametric bootstrap resamples and bias-corrected and accelerated (BCa) 95% confidence intervals.

### Feature-level stability and biological interpretation

To characterize molecular features contributing to stable latent components, we integrated sparse SGCCA weights with loading-based feature characterization. Sparse weights identify conditionally selected features contributing directly to latent component construction, whereas loadings capture correlations between individual features and the latent molecular axis. Because SGCCA enforces sparse conditional selection, correlated features may be excluded despite exhibiting reproducible alignment with the latent component. Feature-level reproducibility was assessed using selection frequency, weight consistency, loading magnitude, and loading sign consistency across repeated cross-validation iterations (**Supplementary Table 8**). Features demonstrating stable loading direction and magnitude were prioritized for downstream interpretation. Pairwise Spearman correlation networks with false discovery rate (FDR) correction characterized cross-omics relationships among all stable metabolomic and inflammation proteomic features. Biological interpretation was performed using joint pathway over-representation analyses applied to stability-filtered feature sets. Two feature sets were evaluated for latent component (LC)1 (M) and LC1(P): features selected directly by the sparse model based on stable weights, and an expanded set additionally including features with stable loadings exceeding the ±0.3 threshold. Joint pathway over-representation analysis was performed using IMPaLA (Integrated Molecular Pathway Level Analysis)^30^, which simultaneously evaluates metabolomic and inflammation proteomic features against integrated pathway databases. Proteomic features were represented by Olink Inflammation panel proteins and metabolomic features by HMDB^31^ (5.0) identifiers. Joint pathway significance was assessed using the IMPaLA joint enrichment p-value, with pathways prioritized when both overlapping inflammation proteins and metabolites were present, reflecting coordinated multi-omics pathway convergence. To visualize pathway-feature relationships, a feature-pathway network was constructed by connecting proteins and metabolites to their overlapping IMPaLA pathways, with Reactome^32^ pathway hierarchy relationships additionally incorporated. The network was visualized in Cytoscape (version 3.10.1)^33^ to illustrate the recovery of coordinated multi-omics pathway structure following inclusion of loading-associated features.

### Statistical analysis

The analytical framework distinguished three distinct sources of variation, each summarized by a different interval. First, the robustness and reproducibility of latent structure across repeated model refits were evaluated through repeated cross-validation procedure; summaries from these analyses (medians, interquartile ranges [IQRs], and directional consistency across resampling iterations) describe the variability of effect estimates across cross-validation repetitions and characterize estimator stability under resampling. These resampling IQRs are not confidence intervals and do not represent population sampling uncertainty. Second, nonparametric subject-level bootstrap resampling (5,000 resamples) quantified sampling uncertainty in effect-size estimates derived from aggregated OOS component scores; mean differences and regression coefficients from these analyses are reported with 95% bias-corrected and accelerated (BCa) confidence intervals, which quantify bootstrap sampling uncertainty for the corresponding effect estimates. Third, where odds ratios are reported from single fitted logistic-regression models for feature-level characterization (**Fig. 5a**), intervals are conventional model-based 95% confidence intervals. All latent component scores, association estimates for the primary disease-association and temporal analyses were derived exclusively from OOS projections. Gardner-Altman estimation plots were generated using dabestr to visualize group differences together with bootstrap-derived sampling distributions. Inference focused on reproducibility, effect-size estimation, and consistency across resampling rather than single-model hypothesis testing, in line with recommendations for estimation-focused statistical practice^29,34–36^.

## Data availability

The plasma inflammation proteins and plasma untargeted annotated metabolites have been published previously^26^. The raw data supporting the current study are available from the corresponding author upon request subject to ethical and legislative review.

## Code Availability

To foster the reproducibility of all the results, the corresponding code implementing SGCCA in repeated cross-validation framework, as well as evaluating the component- and feature-level stability, is available under the MIT license via GitHub at: https://github.com/guanhaibin/stability-driven-multiomics

## Acknowledgements

This work was supported in part by the National Institutes of Health through grants P30ES023515, U2CES030859, R01ES036725, and UL1TR004419.

## Author contributions

H.G. developed the framework, implemented the methodology, performed the analyses, and generated the figures. L.M.P. supervised the project. L.M.P. and M.v.G. acquired funding. H.G. and L.M.P. wrote the initial draft of the manuscript. G.D. contributed to the LC–MS metabolomics data acquisition and quality control. S.K.S. contributed to the Olink data acquisition and quality control. E.C. provided expertise in statistical analysis. All authors contributed to reviewing and editing the manuscript. All authors read and approved the manuscript.

## Additional information

### Supplementary information

Correspondence and requests for materials should be addressed to Haibin Guan.

## Reference

1. India Aldana, S., et al. Pregnancy as a Susceptible Period to Ambient Air Pollution Exposure on the Maternal Postpartum Metabolome. Environ. Sci. Technol. 59, 6400– 6413 (2025).

2. Menyhárt, O. & Gyorffy, B. Multi-omics approaches in cancer research with applications in tumor subtyping, prognosis, and diagnosis. Comput. Struct. Biotechnol. J. 19, 949–960 (2021).

3. Singh, A. et al. DIABLO: an integrative approach for identifying key molecular drivers from multi-omics assays. Bioinformatics 35, 3055–3062 (2019).

4. Tenenhaus, A. et al. Variable selection for generalized canonical correlation analysis. Biostatistics 15, 569–583 (2014).

5. Argelaguet, R. et al. MultiDOmics Factor Analysis—a framework forunsupervised integration of multiDomics data sets. Mol. Syst. Biol. 14, e8124 (2018).

6. Argelaguet, R. et al. MOFA+: a statistical framework for comprehensive integration of multi-modal single-cell data. Genome Biol. 21, (2020).

7. Lock, E. F., Hoadley, K. A., Marron, J. S. & Nobel, A. B. Joint and individual variation explained (JIVE) for integrated analysis of multiple data types. Ann. Appl. Stat. 7, (2013).

8. Zitnik, M. & Zupan, B. Data Fusion by Matrix Factorization. IEEE Trans. Pattern Anal. Mach. Intell. 37, 41–53 (2015).

9. Chalise, P. & Fridley, B. L. Integrative clustering of multi-level ‘omic data based on non-negative matrix factorization algorithm. PLOS ONE 12, e0176278 (2017).

10. Varoquaux, G. Cross-validation failure: Small sample sizes lead to large error bars. NeuroImage 180, 68–77 (2018).

11. Helmer, M. et al. On the stability of canonical correlation analysis and partial least squares with application to brain-behavior associations. *Commun*. Biol. 7, 217 (2024).

12. McCabe, S. D., Lin, D.-Y. & Love, M. I. Consistency and overfitting of multi-omics methods on experimental data. Brief. Bioinform. 21, 1277–1284 (2020).

13. Mihalik, A. et al. Canonical Correlation Analysis and Partial Least Squares for Identifying Brain–Behavior Associations: A Tutorial and a Comparative Study. Biol. Psychiatry Cogn. Neurosci. Neuroimaging 7, 1055–1067 (2022).

14. Rohart, F., Gautier, B., Singh, A. & Lê Cao, K.-A. mixOmics: An R package for ‘omics feature selection and multiple data integration. PLOS Comput. Biol. 13, e1005752 (2017).

15. Dollinger, E. P., Silkwood, K., Atwood, S., Nie, Q. & Lander, A. D. Statistically principled feature selection for single cell transcriptomics. BMC Bioinformatics 26, 238 (2025).

16. Fleiss, J. L. Measuring nominal scale agreement among many raters. Psychol. Bull. 76, 378–382 (1971).

17. Lukaszuk, T., Krawczuk, J., Żyła, K. & Kęsik, J. Stability of Feature Selection in Multi-Omics Data Analysis. Appl. Sci. 14, 11103 (2024).

18. Meinshausen, N. & Bühlmann, P. Stability Selection. J. R. Stat. Soc. Ser. B Stat. Methodol. 72, 417–473 (2010).

19. Hofner, B., Boccuto, L. & Göker, M. Controlling false discoveries in high-dimensional situations: boosting with stability selection. BMC Bioinformatics 16, 144 (2015).

20. Pusa, T. & Rousu, J. Stable biomarker discovery in multi-omics data via canonical correlation analysis. PLOS ONE 19, e0309921 (2024).

21. Hédou, J. et al. Discovery of sparse, reliable omic biomarkers with Stabl. Nat. Biotechnol. 42, 1581–1593 (2024).

22. Muller, E., Shiryan, I. & Borenstein, E. Multi-omic integration of microbiome data for identifying disease-associated modules. Nat. Commun. 15, 2621 (2024).

23. Witten, D. M., Tibshirani, R. & Hastie, T. A penalized matrix decomposition, with applications to sparse principal components and canonical correlation analysis. Biostatistics 10, 515–534 (2009).

24. Tenenhaus, M., Tenenhaus, A. & Groenen, P. J. F. Regularized Generalized Canonical Correlation Analysis: A Framework for Sequential Multiblock Component Methods. Psychometrika 82, 737–777 (2017).

25. Van Gerwen, M. et al. Per- and polyfluoroalkyl substances (PFAS) exposure and thyroid cancer risk. eBioMedicine 97, 104831 (2023).

26. Joseph, G. et al. Inflammatory proteins in pre-diagnosis versus at-diagnosis samples associated with differentiated thyroid cancer. Int. J. Cancer 158, 2072–2082 (2026).

27. Xu, B. et al. Limited generalizability of multivariate brain-based dimensions of child psychiatric symptoms. Commun. Psychol. 2, 16 (2024).

28. Nogueira, S., Sechidis, K. & Brown, G. On the Stability of Feature Selection Algorithms.

29. Ho, J., Tumkaya, T., Aryal, S., Choi, H. & Claridge-Chang, A. Moving beyond P values: data analysis with estimation graphics. Nat. Methods 16, 565–566 (2019).

30. Kamburov, A., Cavill, R., Ebbels, T. M. D., Herwig, R. & Keun, H. C. Integrated pathway-level analysis of transcriptomics and metabolomics data with IMPaLA. Bioinformatics 27, 2917–2918 (2011).

31. Wishart, D. S. et al. HMDB 5.0: the Human Metabolome Database for 2022. Nucleic Acids Res. 50, D622–D631 (2022).

32. Ragueneau, E. et al. The Reactome Knowledgebase 2026. Nucleic Acids Res. 54, D673–D681 (2026).

33. Shannon, P. et al. Cytoscape: A Software Environment for Integrated Models of Biomolecular Interaction Networks. Genome Res. 13, 2498–2504 (2003).

34. Wasserstein, R. L., Schirm, A. L. & Lazar, N. A. Moving to a World Beyond “ *p* 0.05”. Am. Stat. 73, 1–19 (2019).

35. Wasserstein, R. L. & Lazar, N. A. The ASA Statement on *p* -Values: Context, Process, and Purpose. Am. Stat. 70, 129–133 (2016).

36. Calin-Jageman, R. J. & Cumming, G. The New Statistics for Better Science: Ask How Much, How Uncertain, and What Else Is Known. Am. Stat. 73, 271–280 (2019).

