## Supplementary material for "Stability-driven multi-omics integration for reproducible latent structure": SI: Supplementary Information.docx

**Table of Contents**

**Supplementary Method 1:** Repeated cross-validation framework

**Supplementary Method 2:** Metabolomics and inflammatory proteins measurements and preprocessing

**Supplementary Figure 1:** Hyperparameter tuning of SGCCA

**Supplementary Figure 2:** Single-fit SGCCA representation of the cross-omics correlation structure

**Supplementary Table 1:** Summary of component-level stability metrics (score reproducibility and weight stability)

**Supplementary Table 2:** Stratified out-of-sample odds ratios summarized across cross-validation runs (median, interquartile range, and directional consistency)

**Supplementary Figure 3:** Out-of-sample latent component score differences between cases and controls in pre-diagnosis and at-diagnosis samples (n = 162).

**Supplementary Figure 4:** Sensitivity analysis of paired out-of-sample latent component score differences within the pre-diagnosis and at-diagnosis samples in the matched samples (n = 152).

**Supplementary Table 3:** Summary of out-of-sample effect estimates across resampling (median, IQR, and directional consistency)

**Supplementary Figure 5:** Feature-level stability across all latent components

**Supplementary Table 4:** Feature-level stability metrics for the metabolomic latent component (LC1 [M])

**Supplementary Table 5:** Feature-level stability metrics for the inflammatory proteins latent component (LC1 [P])

**Supplementary Figure 6:** Heatmap of pairwise Pearson correlations among inflammatory proteomic features.

**Supplementary Figure 7:** Multi-omics correlation network of inflammatory proteins and metabolites identified by SGCCA.

**Supplementary Table 6:** Joint pathway over-representation analysis results for stable latent component features using IMPaLA.

**Supplementary Table 7:** Baseline characteristics of study population

**Supplementary Table 8**: Stability metrics used to assess latent components and feature-level stability

**Supplementary Method 1: Repeated Cross Validation framework**

We implemented a repeated cross-validation framework to evaluate the stability and out-of-sample performance of sparse generalized canonical correlation analysis (SGCCA).

A repeated K-fold cross-validation scheme was applied with K = 5 folds and R = 200 repetitions on a dataset of n = 162 samples. At each repetition, samples were partitioned into five approximately equal folds, yielding n=130 samples for training (in-sample) and n=32 samples for testing (out-of-sample) per fold. Stratified sampling was used to preserve both outcome class balance (cases vs controls) and temporal structure (e.g., pre-diagnosis vs at-diagnosis populations) within each fold. For each repetition and fold, the training subset was used for preprocessing and model fitting, while the held-out subset was used exclusively for evaluation. For each omics block, features were residualized with respect to covariates using linear models fitted in the training data. The estimated regression coefficients were then applied to both training and test data to obtain residuals. Following residualization, features were standardized using training-based parameters (mean-centering and scaling to unit variance), and the same transformation was applied to the test data. SGCCA models were fitted in the training data using tuned numbers of components and sparsity parameters. The fitted models were then used to project the test data into latent component space, generating out-of-sample component scores.

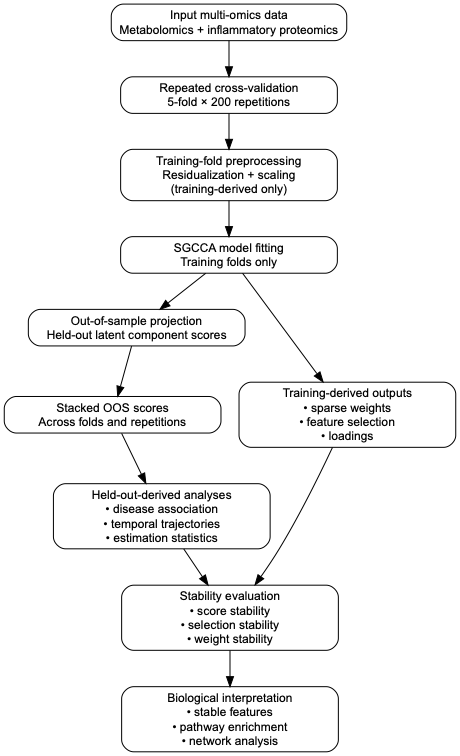
**Supplementary Method 2: Metabolomics and Inflammatory Proteins Measurements and Preprocessing**

**Plasma Metabolomics: Sample Preparation, LC-HRMS Analysis, and Data Preprocessing**

Plasma samples were available for 176 participants for metabolomics analysis. Plasma metabolomics profiling was conducted using liquid chromatography–high resolution mass spectrometry (LC-HRMS) following established protocols^1,2^.

Plasma samples stored at −80 °C were thawed on ice, vortexed, and 35 μL aliquots combined with 105 μL of ice-cold methanol containing internal standards. After incubation at −80 °C for 30 min to precipitate proteins, the samples were centrifuged, and two 50 μL aliquots of supernatant were dried down and stored at -80 °C until analysis. A pooled quality control (QC) sample was generated by combining an additional 5 μL aliquot from each plasma sample. The matrix blank (replacing the plasma with water) and multiple pooled QC samples were extracted and dried following the same protocol. Untargeted metabolomics analysis was performed using LC-HRMS. Prior to analysis, dried extracts were reconstituted in 100% methanol (RPN) or 80% acetonitrile (ZHP), depending on the mode. Samples were analyzed using reverse-phase (RP) and hydrophilic interaction liquid chromatography (HILIC) connected to HRMS in negative (RPN) and positive (ZHP) mode, respectively. Samples were analyzed in a randomized order with pooled QCs injected routinely throughout the run. Metabolite identifications were made using our local database, which comprises over 1000 biologically and environmentally relevant reference standards analyzed under the same conditions. Matching involved considering retention time (± 8 sec), accurate mass (<20ppm), isotope distribution, and MS/MS fragmentation pattern (when available) against the in-house Personal Chemical Database Library and Profinder version B.08.00 Service Pack 3(Agilent Technologies, Santa Clara, USA), resulting in annotation confidence levels 1-3^3^. Metabolite classes were determined using ClassyFire^4^. Annotated metabolites in RPN and ZHP were filtered and normalized, respectively. Metabolites with more than 20% missing values or a signal-to-noise ratio less than 1.5-fold compared to solvent blanks were excluded. Batch effects were corrected using the Quality Control Robust Spline Correction (QCRSC)^5^ method. A quality control criterion of 30% coefficient of variation (CV) across repeated injections of pooled QC samples was used to remove metabolites with high variability. The filtered and normalized metabolites from both modes were then combined, with any duplicates removed; priority was given to the mode with higher abundance and fewer missing values. A log2 transformation was applied to the remaining metabolites to normalize the intensity. Missing values were then imputed using the random forest algorithm from R package ("MissForest")^6^. A total of 284 annotated plasma metabolites were retained for downstream analyses.

**Inflammatory Proteomics: Olink Assay and Data Preprocessing**

Inflammatory protein profiling was performed on the same plasma samples using the Olink Target 96 Inflammation panel, which quantifies 92 inflammation-related proteins simultaneously. The assay is based on Proximity Extension Assay (PEA) technology, a dual-recognition immunoassay coupled with quantitative polymerase chain reaction (qPCR) readout. Briefly, each target protein is recognized by a pair of oligonucleotide-labeled antibodies that bind to distinct epitopes. Upon simultaneous binding to the target protein, the oligonucleotide probes are brought into proximity, allowing hybridization and enzymatic extension to form a unique DNA barcode. This sequence is subsequently amplified using real-time PCR, enabling sensitive and specific detection of protein abundance. Following incubation and extension, samples were processed using the Biomark HD System (Fluidigm) for high-throughput qPCR detection. The resulting data are reported as Normalized Protein eXpression (NPX) values, which are on a log2 scale and represent relative protein abundance after normalization. Quality control procedures included the use of internal assay controls added to each sample to monitor key steps of the workflow, including immunoreaction, extension, and amplification/detection. In addition, inter-plate controls and negative controls were used to assess technical variation and background signal. Data normalization was performed using Olink’s standard normalization procedures, which account for both intra- and inter-plate variation. Proteins with values below the limit of detection (LOD), as defined by Olink, were flagged, and proteins with a high proportion of measurements below LOD were excluded from downstream analyses where appropriate. A total of 92 plasma inflammatory proteins were included for the multi-omics data integration.

Inflammatory protein profiling was conducted in 166 plasma samples with sufficient volume. Sample-level quality control was performed according to the manufacturer’s guidelines by assessing deviation from the median of internal controls. Samples with a deviation of less than 0.3 NPX from the median were considered to pass quality control. Of the 166 samples analyzed, 162 met the quality control criteria and were included in downstream analyses. Of those 162 available samples, 152 of them remain complete paired matched case controls.

**Supplementary Figure 1: Hyperparameter tuning of SGCCA**

SGCCA hyperparameters were selected using the same repeated cross-validation framework. Sparsity parameters were first optimized with the number of components fixed to one per block. Subsequently, the number of components was tuned while holding the selected sparsity values fixed. Both steps used 5-fold cross-validation repeated 200 times, consistent with the main analysis. Predictive performance was assessed using balanced accuracy, based on a regularized logistic regression model fitted on the derived latent components.
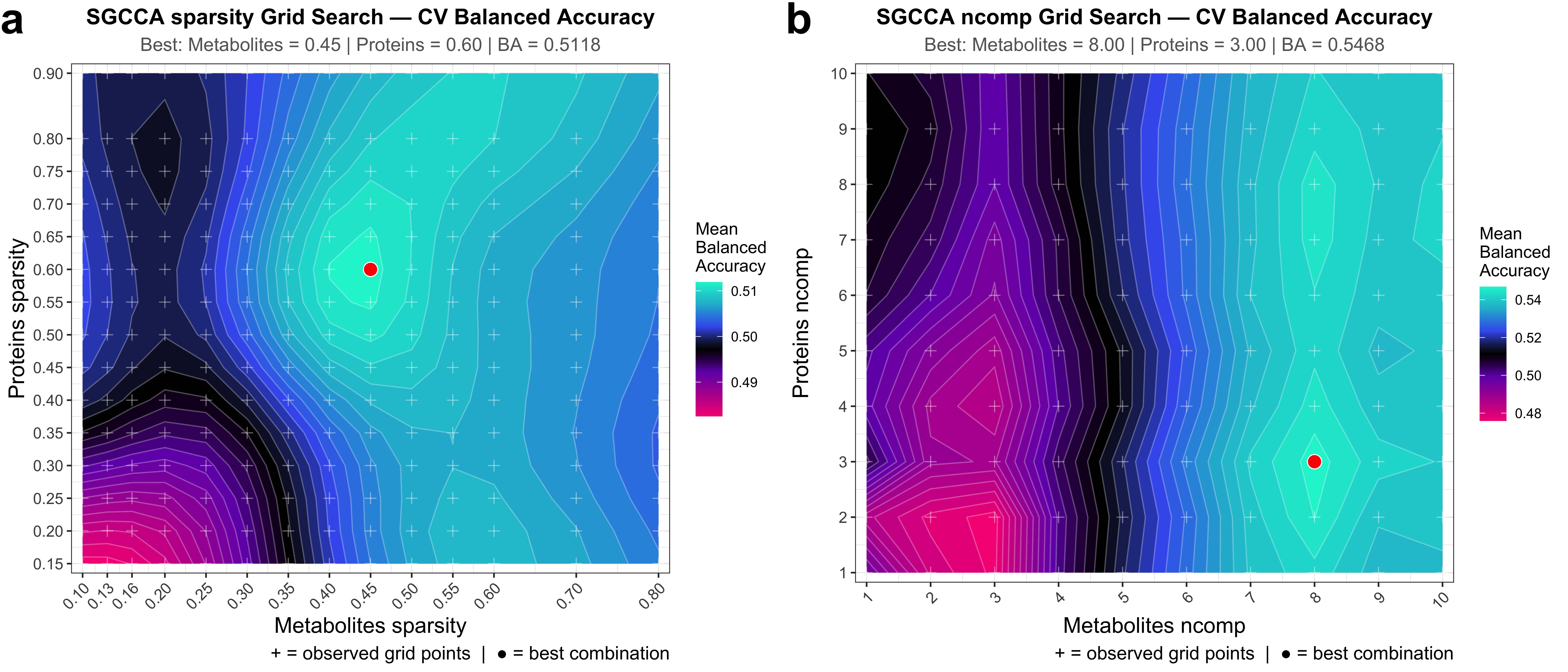

**Supplementary Figure 2: Single-fit SGCCA representation of cross-omics correlation structure**

A representative single-fit SGCCA model applied to the metabolomics and proteomics blocks, showing the cross-omics structure captured by s ingle model fit. a. Feature-level correlation structure within and between omics blocks. b. Correlation heatmap of latent components derived from SGCCA.

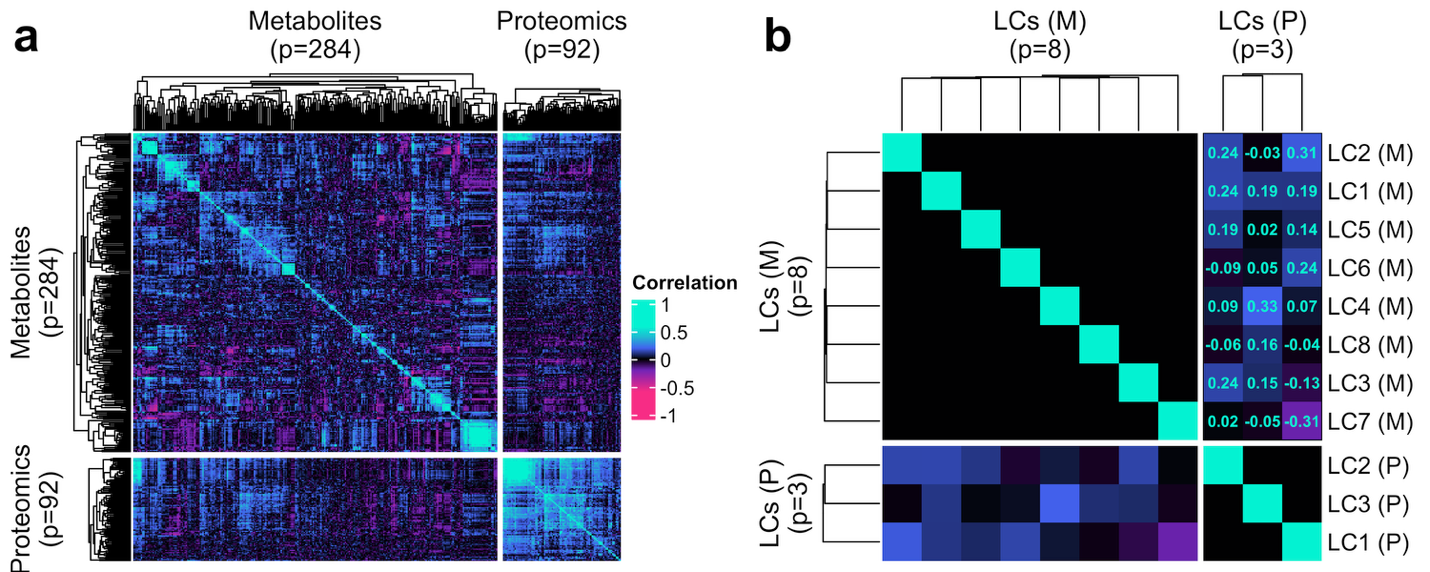

**Supplementary Table 1:** Summary of component-level stability metrics

Selection stability (selection rate, Nogueira index, and Jaccard index) and component stability metrics are summarized across resampling. Component stability is quantified using pairwise Spearman’s rho of out-of-sample scores and feature weights, reported as median [Q25–Q75].

| **Block** | **LC** | **Selection Rate (%)** | **Nogueira Index** | **Jaccard Index** | **Weight**  **Median [Q25, Q75]** | **Score**  **Median [Q25, Q75]** |
| --- | --- | --- | --- | --- | --- | --- |
| **Metabolomics** | *LC1 (M)* | *27.1 [22.2–33.8]* | *0.40* | *0.39* | *0.67 [0.64, 0.71]* | *0.59 [0.58, 0.60]* |
|  | *LC2 (M)* | *26.1 [21.1–34.2]* | *0.27* | *0.31* | *0.44 [0.34, 0.52]* | *0.47 [0.45, 0.49]* |
|  | *LC3 (M)* | *26.1 [21.5–33.1]* | *0.14* | *0.23* | *0.42 [0.35, 0.48]* | *0.24 [0.22, 0.26]* |
|  | *LC4 (M)* | *26.4 [21.5–33.5]* | *0.09* | *0.20* | *0.32 [0.26, 0.38]* | *0.23 [0.20, 0.25]* |
|  | *LC5 (M)* | *26.8 [21.5–33.5]* | *0.05* | *0.18* | *0.18 [0.12, 0.24]* | *0.18 [0.16, 0.21]* |
|  | *LC6 (M)* | *26.8 [21.8–35.2]* | *0.04* | *0.18* | *0.22 [0.15, 0.28]* | *0.17 [0.15, 0.20]* |
|  | *LC7 (M)* | *27.1 [22.2–34.9]* | *0.02* | *0.17* | *0.18 [0.12, 0.24]* | *0.11 [0.09, 0.13]* |
|  | *LC8 (M)* | *27.1 [20.8–34.2]* | *0.03* | *0.17* | *0.15 [0.09, 0.21]* | *0.09 [0.07, 0.12]* |
| **Inflammatory**  **Proteomics** | *LC1 (P)* | *57.6 [45.7–73.9]* | *0.37* | *0.59* | *0.91 [0.88, 0.93]* | *0.76 [0.75, 0.77]* |
|  | *LC2 (P)* | *54.3 [45.7–69.6]* | *0.52* | *0.65* | *0.76 [0.73, 0.79]* | *0.69 [0.67, 0.72]* |
|  | *LC3 (P)* | *58.7 [47.8–72.8]* | *0.12* | *0.47* | *0.37 [0.31, 0.42]* | *0.50 [0.47, 0.54]* |

**Note:** Selection rate (%) is reported as median (min–max) across resampling iterations. The Nogueira index accounts for chance agreement under sparsity and is the primary measure of selection stability, whereas the Jaccard index is provided for comparison without sparsity correction. For interpretability, commonly used reference ranges are shown (e.g., Nogueira index >0.5, 0.25–0.5, <0.25), but these are intended as descriptive guidelines rather than strict thresholds. Score stability reflects reproducibility of out-of-sample latent component scores, and weight stability reflects concordance of feature weight rankings across resampling.

**Supplementary Table 2:** Stratified out-of-sample odds ratios summarized across cross-validation runs (median, interquartile range, and directional consistency).

Stratified association estimates were derived from repeated cross-validation analyses. Reported ORs represent the median out-of-sample estimate across validation runs, summarized by the interquartile range (IQR; 25th–75th percentiles) of the out-of-sample estimate distribution across runs. These summaries describe the variability of effect estimates across cross-validation repetitions and characterize estimator stability rather than population sampling uncertainty. The proportions of positive and negative estimates correspond to the fraction of runs yielding OR > 1 and OR < 1, respectively.

| **LC** | **Group** | **Median OR [IQR]** | **% positive** | **% negative** |
| --- | --- | --- | --- | --- |
| **LC1 (M)** | *At-Diagnosis* | *0.93 [0.85, 1.02]* | *31.5%* | *68.5%* |
|  | *Pre-Diagnosis* | *0.73 [0.65, 0.82]* | *3.5%* | *96.5%* |
|  | *Total* | *0.86 [0.80, 0.92]* | *7.5%* | *92.5%* |
| **LC1 (P)** | *At-Diagnosis* | *1.07 [1.03, 1.11]* | *86.0%* | *14.0%* |
|  | *Pre-Diagnosis* | *1.85 [1.67, 1.98]* | *99.0%* | *1.0%* |
|  | *Total* | *1.23 [1.18, 1.28]* | *96.0%* | *4.0%* |
| **LC2 (P)** | *At-Diagnosis* | *0.88 [0.81, 0.96]* | *16.5%* | *83.5%* |
|  | *Pre-Diagnosis* | *1.12 [1.02, 1.20]* | *81.0%* | *19.0%* |
|  | *Total* | *0.97 [0.91, 1.04]* | *40.5%* | *59.5%* |

**Supplementary Figure 3:** Out-of-sample latent component score differences between cases and controls in pre-diagnosis and at-diagnosis samples (n = 162).

Estimation plots comparing subject-level averaged out-of-sample (OOS) latent component scores between cases and controls within the pre-diagnosis and at-diagnosis samples in the full dataset (n = 162). Points represent individual samples. Mean differences (At-diagnosis − Pre-diagnosis) are shown with bias-corrected and accelerated (BCa) 95% bootstrap confidence intervals derived from 5,000 subject-level bootstrap resamples.

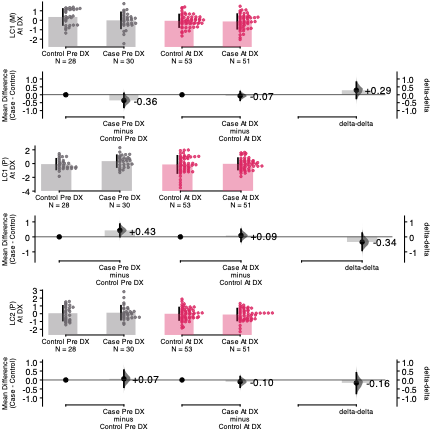

**Supplementary Figure 4:** Sensitivity analysis of paired out-of-sample latent component score differences within the pre-diagnosis and at-diagnosis samples in the matched samples (n = 152).

Mean paired differences are shown with bias-corrected and accelerated (BCa) 95% bootstrap confidence intervals derived from 5,000 subject-level bootstrap resamples.

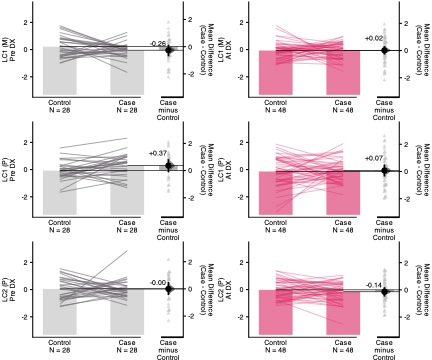

**Supplementary Table 3:** Estimation-statistics summary of OOS component score differences between Pre-diagnosis and At-diagnosis groups.

Mean differences were calculated as At-diagnosis minus Pre-diagnosis using subject-level averaged OOS scores. Confidence intervals represent 95% bias-corrected and accelerated bootstrap intervals from 5,000 subject-level resamples. Permutation t-test p-values are reported. Meanwhile, repeated cross-validation stability of components score and the time to diagnosis/reference relationship was summarized as the median, interquartile range, and directional consistency of the effect estimates across 200 CV repetitions.

|  | | **At-diagnosis vs Pre-diagnosis** | | **Time to diagnosis / reference** | | |
| --- | --- | --- | --- | --- | --- | --- |
| **LC** | ***Group*** | ***Mean difference (95% CI)*** | ***p-value*** | ***Median slope (IQR)*** | ***% positive*** | ***% negative*** |
| LC1 (M) | *Cases* | *-0.08 (-0.45, 0.28)* | *0.659* | *0.01 (-0.02, 0.03)* | *56.0%* | *44.0%* |
|  | *Controls* | *-0.38 (-0.75, -0.02)* | *0.030* | *-0.04 (-0.06, -0.03)* | *3.5%* | *96.5%* |
| LC1 (P) | *Cases* | *-0.40 (-0.78, -0.003)* | *0.045* | *-0.10 (-0.11, -0.09)* | *0.0%* | *100.0%* |
|  | *Controls* | *-0.05 (-0.51, 0.37)* | *0.849* | *-0.01 (-0.02, 0.001)* | *28.5%* | *71.5%* |
| LC2 (P) | *Cases* | *-0.24 (-0.69, 0.12)* | *0.228* | *-0.05 (-0.06, -0.03)* | *4.0%* | *96.0%* |
|  | *Controls* | *-0.07 (-0.51, 0.34)* | *0.713* | *-0.01 (-0.02, 0.01)* | *30.0%* | *70.0%* |

**Note:** Mean differences are computed as At-diagnosis minus Pre-diagnosis (negative values indicate higher scores in the pre-diagnostic group). Slopes represent the change in LC score per unit increase in time to diagnosis/reference (years); negative slopes indicate higher scores further from diagnosis, i.e., earlier in the pre-diagnostic window. Directional consistency (% positive / % negative) is computed across 200 CV repetitions.

**Supplementary Figure 5: Feature-level stability across all derived latent components**

a. Forest plots and loading patterns for selected stable features from metabolomics (left) and proteomics (right) blocks. Odds ratios (ORs) with 95% confidence intervals were estimated using logistic regression adjusted for age, sex, BMI, race/ethnicity and sample storage time. Corresponding loading values across latent components (LCs) are shown as heatmaps, with color indicating the magnitude and direction of the mean loading. Features with stable weights for each LC are highlighted. The stacked bar plot summarizes the number of metabolites colored by superclass contributing to individual components. b. Scatter plots show median feature weights (x-axis) versus mean loadings (y-axis) for each LC in metabolomics (top) and proteomics (bottom). Each dot represents a feature, colored by the combined index (sum of median weight and mean loading), reflecting concordance between weight and loading. Vertical and horizontal reference lines indicate zero weight and loading, respectively. Dashed horizontal lines at ±0.3 are shown as reference guides to indicate features with relatively larger loading magnitudes.

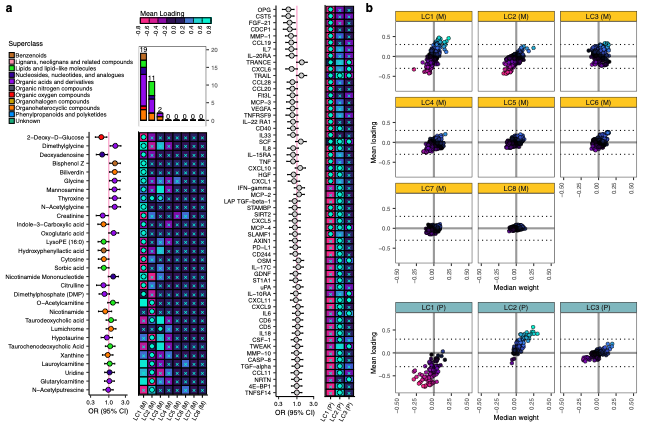

**Supplementary Table 4:** **Feature-level stability metrics for the metabolomic latent component (LC1 (M))**

For each feature, selection frequency, distribution of non-zero sparse weights (median [Q25–Q75]), and loading stability metrics (mean ± SD and sign consistency) across resampling are reported. These metrics quantify the reproducibility and directional consistency of feature contributions to the LC1 (M).

|  | **Weight** | | **Loading** | |
| --- | --- | --- | --- | --- |
| **Metabolite** | ***Selection Frequency*** | ***Median [Q25, Q75]*** | ***Sign Consistency*** | ***Mean ± SD*** |
| 2-Deoxy-D-Glucose | *0.999* | *-0.287 [-0.338, -0.233]* | *0.999* | *-0.322 ± 0.113* |
| Dimethylglycine | *0.994* | *0.208 [0.16, 0.261]* | *0.999* | *0.456 ± 0.157* |
| Biliverdin | *0.979* | *0.177 [0.121, 0.232]* | *0.998* | *0.377 ± 0.117* |
| Thyroxine | *0.971* | *0.17 [0.112, 0.236]* | *0.881* | *0.111 ± 0.096* |
| Glycine | *0.952* | *0.162 [0.112, 0.218]* | *0.965* | *0.287 ± 0.142* |
| Deoxyadenosine | *0.964* | *-0.158 [-0.215, -0.103]* | *0.882* | *-0.15 ± 0.122* |
| Oxoglutaric acid | *0.943* | *0.151 [0.097, 0.205]* | *0.993* | *0.252 ± 0.109* |
| Mannosamine | *0.947* | *0.149 [0.094, 0.209]* | *0.831* | *0.178 ± 0.176* |
| Bisphenol Z | *0.942* | *0.149 [0.097, 0.205]* | *1.000* | *0.323 ± 0.091* |
| N-Acetylglycine | *0.958* | *0.148 [0.092, 0.206]* | *1.000* | *0.363 ± 0.079* |
| Creatinine | *0.931* | *-0.142 [-0.19, -0.095]* | *0.998* | *-0.389 ± 0.137* |
| Cytosine | *0.900* | *-0.115 [-0.168, -0.066]* | *1.000* | *-0.373 ± 0.09* |
| Dimethylphosphate (DMP) | *0.799* | *-0.109 [-0.172, -0.053]* | *0.674* | *-0.085 ± 0.149* |
| Indole-3-Carboxylic acid | *0.846* | *-0.107 [-0.163, -0.059]* | *0.992* | *-0.203 ± 0.1* |
| LysoPE (16:0) | *0.885* | *-0.104 [-0.157, -0.058]* | *0.992* | *-0.274 ± 0.137* |
| Hydroxyphenyllactic acid | *0.847* | *-0.103 [-0.155, -0.061]* | *0.999* | *-0.272 ± 0.114* |
| Sorbic acid | *0.830* | *-0.098 [-0.15, -0.054]* | *1.000* | *-0.372 ± 0.119* |
| Citrulline | *0.851* | *-0.097 [-0.149, -0.053]* | *1.000* | *-0.343 ± 0.11* |
| Nicotinamide Mononucleotide | *0.804* | *0.096 [0.049, 0.146]* | *0.992* | *0.35 ± 0.156* |
| LysoPC (14:0) | *0.793* | *-0.094 [-0.149, -0.051]* | *0.946* | *-0.2 ± 0.139* |
| Glycocholic acid | *0.819* | *-0.093 [-0.153, -0.051]* | *0.995* | *-0.279 ± 0.112* |
| Bilirubin | *0.717* | *0.093 [0.048, 0.143]* | *0.971* | *0.274 ± 0.134* |
| Hydroxykynurenine | *0.733* | *-0.091 [-0.139, -0.047]* | *0.998* | *-0.295 ± 0.1* |
| N-Acetylneuraminic acid | *0.703* | *-0.09 [-0.142, -0.044]* | *0.530* | *-0.003 ± 0.119* |
| Asymmetrical dimethylarginine | *0.798* | *0.088 [0.044, 0.14]* | *0.990* | *0.439 ± 0.185* |
| 4-Hydroxybenzoic acid | *0.722* | *-0.087 [-0.142, -0.044]* | *0.989* | *-0.163 ± 0.08* |
| cis-Vaccenic Acid / Elaidate/ Oleic acid / Petroselinic acid | *0.798* | *0.086 [0.042, 0.132]* | *0.990* | *0.363 ± 0.196* |
| Mono-n-butylphthalate (BzBP, DnBP) | *0.739* | *0.085 [0.045, 0.14]* | *0.999* | *0.22 ± 0.079* |
| Phenylacetic acid | *0.790* | *-0.084 [-0.135, -0.044]* | *0.998* | *-0.272 ± 0.11* |
| Dipalmitoyl-Phosphoethanolamine | *0.722* | *0.084 [0.044, 0.136]* | *0.950* | *0.126 ± 0.079* |
| Tyrosine /4-Hydroxy-4-(3-pyridyl) butanoic acid | *0.757* | *-0.074 [-0.114, -0.04]* | *0.999* | *-0.425 ± 0.147* |
| N-Palmitoyl Taurine | *0.712* | *0.074 [0.042, 0.121]* | *0.971* | *0.257 ± 0.154* |
| Methyl Vanillate | *0.758* | *-0.073 [-0.113, -0.037]* | *0.999* | *-0.416 ± 0.147* |
| O-Acetylcarnitine | *0.688* | *0.067 [0.034, 0.111]* | *1.000* | *0.471 ± 0.129* |
| Docosahexaenoic acid | *0.612* | *0.066 [0.03, 0.114]* | *0.986* | *0.309 ± 0.172* |
| Aminocaproic acid | *0.674* | *-0.065 [-0.105, -0.03]* | *0.997* | *-0.371 ± 0.14* |
| N-Acetylaspartic acid | *0.562* | *0.063 [0.024, 0.11]* | *0.996* | *0.42 ± 0.156* |
| Hydroxybenzaldehyde | *0.641* | *-0.061 [-0.1, -0.03]* | *0.999* | *-0.413 ± 0.145* |
| 5-Hydroxylysine | *0.531* | *0.06 [0.029, 0.102]* | *0.986* | *0.37 ± 0.177* |
| Glucose | *0.477* | *-0.058 [-0.102, -0.03]* | *0.994* | *-0.372 ± 0.156* |
| Stearic acid | *0.525* | *0.057 [0.026, 0.097]* | *0.971* | *0.317 ± 0.194* |
| Alanine | *0.523* | *-0.057 [-0.104, -0.029]* | *1.000* | *-0.365 ± 0.109* |
| Cystine | *0.547* | *0.057 [0.025, 0.101]* | *0.995* | *0.435 ± 0.18* |
| 4-Hydroxybenzophenone | *0.484* | *-0.053 [-0.092, -0.024]* | *1.000* | *-0.332 ± 0.11* |
| Lysine | *0.338* | *0.046 [0.023, 0.077]* | *0.974* | *0.358 ± 0.191* |
| Palmitic acid | *0.329* | *0.046 [0.02, 0.072]* | *0.974* | *0.323 ± 0.202* |
| Palmitoleic acid | *0.331* | *0.046 [0.021, 0.07]* | *0.983* | *0.321 ± 0.187* |
| N,N,N-Trimethyllysine | *0.298* | *0.041 [0.022, 0.074]* | *0.978* | *0.377 ± 0.188* |
| Myristic acid | *0.268* | *0.039 [0.018, 0.076]* | *0.991* | *0.322 ± 0.181* |
| Octanoylcarnitine | *0.267* | *0.039 [0.018, 0.072]* | *1.000* | *0.331 ± 0.118* |
| 2-Aminoisobutyric acid | *0.142* | *0.038 [0.013, 0.073]* | *0.987* | *0.31 ± 0.125* |
| Adrenic Acid | *0.207* | *0.036 [0.016, 0.07]* | *0.964* | *0.301 ± 0.195* |
| Hexadecanol | *0.231* | *0.031 [0.016, 0.056]* | *0.984* | *0.329 ± 0.186* |
| Heptadecanoic acid | *0.115* | *0.028 [0.014, 0.052]* | *0.975* | *0.308 ± 0.184* |
| Lauroylcarnitine | *0.030* | *0.025 [0.009, 0.053]* | *0.991* | *0.345 ± 0.153* |
| N-Acetylputrescine | *0.041* | *-0.021 [-0.046, -0.007]* | *0.996* | *-0.301 ± 0.116* |
| Glutarylcarnitine | *0.013* | *0.014 [-0.019, 0.024]* | *0.969* | *0.308 ± 0.161* |

**Supplementary Table 5:** **Feature-level stability metrics for the metabolomic latent component (LC1 (P))**

For each feature, selection frequency, distribution of non-zero sparse weights (median [Q25–Q75]), and loading stability metrics (mean ± SD and sign consistency) across resampling are reported. These metrics quantify the reproducibility and directional consistency of feature contributions to the LC1 (P).

|  | **Weight** | | **Loading** | |
| --- | --- | --- | --- | --- |
| **Proteins** | **Selection Frequency** | **Median [Q25, Q75]** | **Sign Consistency** | **Mean ± SD** |
| OPG | 1.000 | -0.288 [-0.323, -0.25] | 0.991 | -0.576 ± 0.098 |
| CST5 | 1.000 | -0.284 [-0.33, -0.241] | 0.991 | -0.408 ± 0.082 |
| FGF-21 | 1.000 | -0.266 [-0.314, -0.221] | 0.992 | -0.383 ± 0.083 |
| CDCP1 | 1.000 | -0.249 [-0.29, -0.205] | 0.991 | -0.522 ± 0.11 |
| MMP-1 | 1.000 | -0.214 [-0.249, -0.175] | 0.991 | -0.668 ± 0.137 |
| CCL19 | 0.996 | -0.2 [-0.242, -0.157] | 0.991 | -0.467 ± 0.103 |
| CXCL6 | 1.000 | -0.199 [-0.235, -0.161] | 0.991 | -0.65 ± 0.151 |
| IL7 | 0.998 | -0.187 [-0.226, -0.152] | 0.991 | -0.635 ± 0.143 |
| IL-20RA | 0.993 | -0.165 [-0.213, -0.123] | 0.991 | -0.485 ± 0.111 |
| MCP-3 | 0.995 | -0.162 [-0.207, -0.122] | 0.991 | -0.586 ± 0.126 |
| TRAIL | 0.961 | 0.157 [0.1, 0.216] | 0.849 | -0.151 ± 0.155 |
| VEGFA | 0.992 | -0.155 [-0.197, -0.119] | 0.991 | -0.722 ± 0.157 |
| CCL20 | 0.986 | -0.15 [-0.193, -0.107] | 0.991 | -0.536 ± 0.112 |
| TNFRSF9 | 0.973 | -0.137 [-0.174, -0.091] | 0.991 | -0.521 ± 0.151 |
| CCL28 | 0.976 | -0.127 [-0.171, -0.09] | 0.991 | -0.543 ± 0.109 |
| TRANCE | 0.929 | 0.126 [0.076, 0.177] | 0.543 | 0.029 ± 0.109 |
| CD40 | 0.982 | -0.125 [-0.159, -0.092] | 0.991 | -0.736 ± 0.162 |
| Flt3L | 0.953 | -0.125 [-0.172, -0.079] | 0.990 | -0.291 ± 0.082 |
| IL8 | 0.962 | -0.12 [-0.151, -0.083] | 0.991 | -0.712 ± 0.144 |
| IL33 | 0.944 | -0.116 [-0.157, -0.072] | 0.991 | -0.426 ± 0.089 |
| IL-15RA | 0.965 | -0.116 [-0.154, -0.077] | 0.991 | -0.572 ± 0.144 |
| TNF | 0.961 | -0.114 [-0.145, -0.08] | 0.991 | -0.554 ± 0.127 |
| SCF | 0.876 | 0.106 [0.063, 0.161] | 0.787 | -0.069 ± 0.109 |
| CXCL10 | 0.814 | 0.096 [0.053, 0.148] | 0.953 | -0.268 ± 0.141 |
| IL-22 RA1 | 0.901 | -0.09 [-0.125, -0.055] | 0.991 | -0.519 ± 0.104 |
| HGF | 0.906 | -0.083 [-0.12, -0.052] | 0.991 | -0.677 ± 0.151 |
| CXCL1 | 0.889 | -0.082 [-0.114, -0.048] | 0.991 | -0.611 ± 0.148 |
| SLAMF1 | 0.810 | -0.076 [-0.113, -0.041] | 0.991 | -0.536 ± 0.13 |
| IL2 | 0.787 | -0.068 [-0.111, -0.033] | 0.991 | -0.362 ± 0.094 |
| MCP-4 | 0.829 | -0.066 [-0.099, -0.039] | 0.991 | -0.588 ± 0.151 |
| IL4 | 0.720 | -0.065 [-0.11, -0.033] | 0.991 | -0.196 ± 0.072 |
| MCP-2 | 0.542 | 0.063 [0.029, 0.106] | 0.988 | -0.406 ± 0.135 |
| IL-12B | 0.664 | -0.062 [-0.102, -0.031] | 0.989 | -0.358 ± 0.121 |
| STAMBP | 0.803 | -0.06 [-0.09, -0.033] | 0.991 | -0.696 ± 0.168 |
| SIRT2 | 0.800 | -0.058 [-0.087, -0.033] | 0.991 | -0.664 ± 0.164 |
| CXCL5 | 0.714 | -0.058 [-0.087, -0.03] | 0.991 | -0.596 ± 0.144 |
| TSLP | 0.668 | -0.055 [-0.084, -0.027] | 0.991 | -0.459 ± 0.101 |
| AXIN1 | 0.730 | -0.055 [-0.086, -0.028] | 0.991 | -0.644 ± 0.16 |
| uPA | 0.396 | 0.055 [0.026, 0.097] | 0.984 | -0.334 ± 0.123 |
| CCL3 | 0.669 | -0.053 [-0.1, -0.027] | 0.991 | -0.469 ± 0.114 |
| LAP TGF-beta-1 | 0.733 | -0.053 [-0.08, -0.029] | 0.991 | -0.702 ± 0.166 |
| LIF-R | 0.650 | -0.053 [-0.088, -0.024] | 0.988 | -0.37 ± 0.119 |
| ADA | 0.623 | -0.05 [-0.081, -0.026] | 0.991 | -0.464 ± 0.114 |
| IL-17A | 0.480 | -0.047 [-0.105, -0.019] | 0.991 | -0.387 ± 0.104 |
| IL10 | 0.385 | -0.045 [-0.08, -0.017] | 0.987 | -0.336 ± 0.112 |
| MCP-1 | 0.478 | -0.044 [-0.081, -0.018] | 0.991 | -0.452 ± 0.114 |
| IL-17C | 0.336 | 0.042 [0.015, 0.079] | 0.988 | -0.412 ± 0.124 |
| CCL4 | 0.330 | -0.038 [-0.071, -0.015] | 0.991 | -0.458 ± 0.12 |
| CD5 | 0.240 | 0.037 [0.013, 0.079] | 0.985 | -0.438 ± 0.144 |
| CSF-1 | 0.334 | -0.036 [-0.062, -0.013] | 0.991 | -0.47 ± 0.127 |
| CD244 | 0.477 | -0.035 [-0.059, -0.016] | 0.991 | -0.649 ± 0.158 |
| PD-L1 | 0.512 | -0.035 [-0.06, -0.017] | 0.991 | -0.677 ± 0.164 |
| IL-18R1 | 0.401 | -0.034 [-0.059, -0.013] | 0.989 | -0.347 ± 0.106 |
| GDNF | 0.351 | -0.033 [-0.062, -0.013] | 0.991 | -0.482 ± 0.116 |
| CXCL11 | 0.457 | -0.032 [-0.055, -0.013] | 0.991 | -0.622 ± 0.16 |
| CXCL9 | 0.279 | 0.032 [0.006, 0.067] | 0.986 | -0.468 ± 0.151 |
| IL6 | 0.241 | 0.032 [0.011, 0.066] | 0.990 | -0.357 ± 0.102 |
| IL-10RB | 0.244 | -0.03 [-0.067, -0.009] | 0.990 | -0.362 ± 0.117 |
| CASP-8 | 0.213 | 0.028 [0.012, 0.064] | 0.991 | -0.629 ± 0.161 |
| FGF-23 | 0.285 | -0.027 [-0.05, -0.006] | 0.988 | -0.35 ± 0.122 |
| ST1A1 | 0.332 | -0.026 [-0.046, -0.009] | 0.991 | -0.521 ± 0.132 |
| FGF-5 | 0.257 | -0.023 [-0.068, 0.013] | 0.989 | -0.334 ± 0.116 |
| 4E-BP1 | 0.277 | -0.019 [-0.042, -0.006] | 0.991 | -0.487 ± 0.129 |
| CCL23 | 0.219 | -0.013 [-0.036, 0.004] | 0.990 | -0.329 ± 0.097 |
| CCL11 | 0.200 | -0.012 [-0.032, 0.005] | 0.991 | -0.413 ± 0.116 |
| TGF-alpha | 0.229 | 0.004 [-0.021, 0.034] | 0.991 | -0.491 ± 0.125 |
| TNFSF14 | 0.187 | -0.004 [-0.022, 0.018] | 0.991 | -0.647 ± 0.163 |
| CX3CL1 | 0.202 | 0.004 [-0.02, 0.044] | 0.975 | -0.339 ± 0.133 |
| NRTN | 0.211 | -0.003 [-0.027, 0.029] | 0.991 | -0.372 ± 0.086 |
| IL18 | 0.201 | 0.003 [-0.021, 0.034] | 0.988 | -0.405 ± 0.129 |
| MMP-10 | 0.186 | 0 [-0.024, 0.022] | 0.991 | -0.399 ± 0.104 |

**Supplementary Figure 6:** Heatmap of pairwise Pearson correlations among inflammatory proteomic features. Rows and columns represent proteins from the inflammatory panel, and are grouped into LC 1 (P) selected features (non-zero sparse weights), high-loading features (stable loadings but not selected) and low-loading features. Rows and columns are split by feature group, with hierarchical clustering performed within each group. Colors indicate pairwise correlation coefficients, ranging from negative (pink) to positive (blue) associations. Selected features show relative stronger and more structured correlations with high-loading features compared to low-loading or other proteins, highlighting a broader network of coordinated molecular variation beyond sparse feature selection.

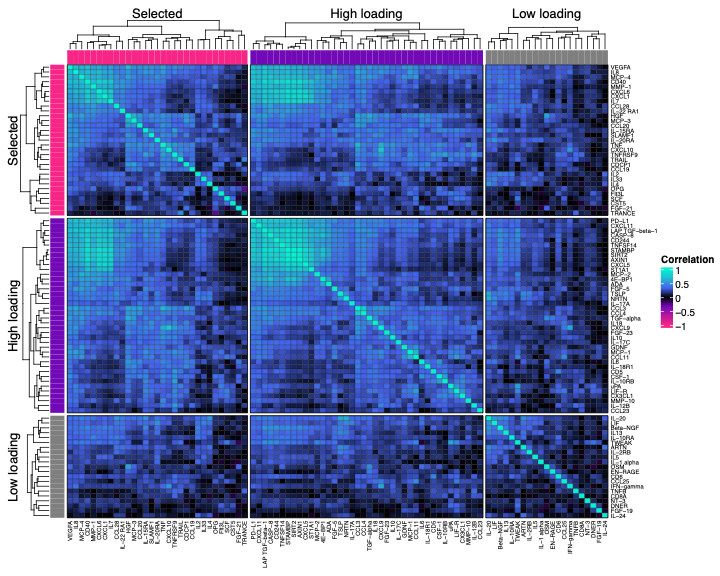

**Supplementary Figure 7:** Multi-omics correlation network of inflammatory proteins and metabolites identified by SGCCA.

A network was constructed to represent correlations between selected inflammatory proteins and metabolites identified through sparse generalized canonical correlation analysis (SGCCA). Pairwise spearman correlations were computed between selected features using residualized values after adjustment for age, sex, race/ethnicity, BMI and sample storage time. The resulting network was visualized using Metscape^11^ and NetworkAnalyzer^12^ in Cytoscape 3.10.1^13^. Nodes represent individual features, including inflammatory proteins and metabolites, and edges represent pairwise Spearman correlations coefficients. Node color indicates molecular type (metabolites vs inflammatory proteins), while node shape represents features selected based on stable SGCCA weights or those additionally identified based on stable loadings. Edges are colored according to the direction of the correlation (blue, positive; magenta, negative), and only significant associations passing multiple testing correction (false discovery rate adjusted p < 0.05) are shown. Node size reflects network centrality, represented by betweenness centrality as calculated using network analysis. In particular, betweenness centrality quantifies the extent to which a node lies on the shortest paths between other nodes, thereby highlighting features that are highly connected and may act as key intermediates linking metabolic and inflammatory processes. This cross-omics network highlights coordinated relationships between metabolic and inflammatory features and reveals potential bridge nodes that may play central roles in mediating interactions between these biological systems.

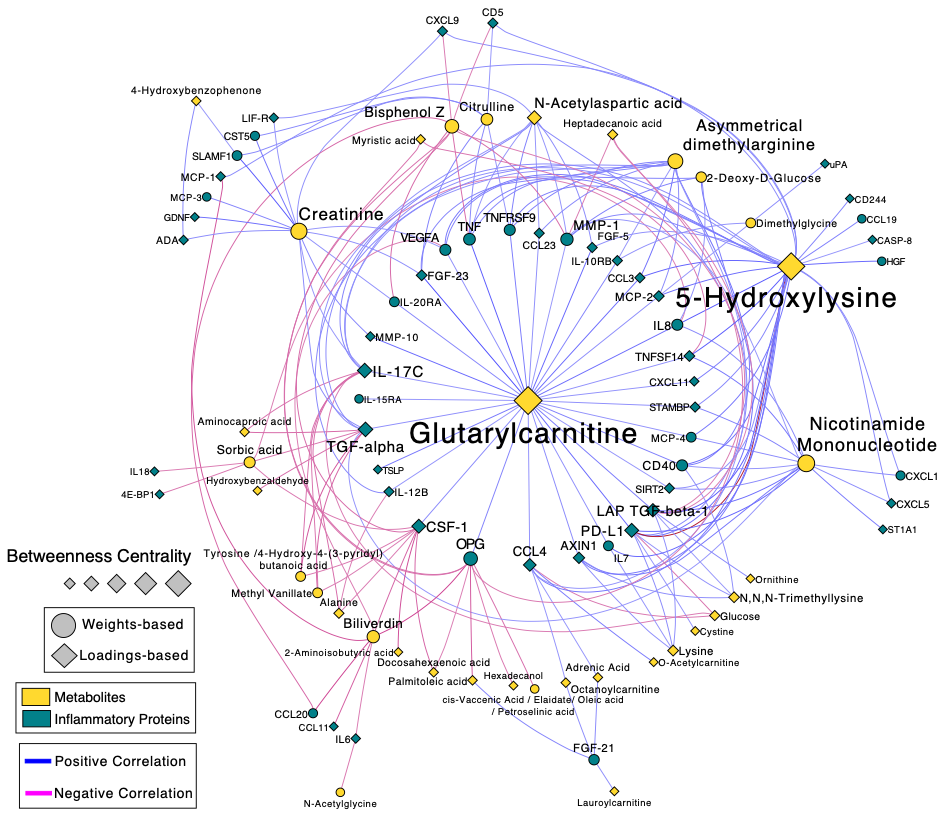

**Supplementary Table 6:** Joint pathway over-representation analysis results for stable latent component features using IMPaLA.

Joint pathway over-representation analysis was performed using IMPaLA against the Reactome pathway database, separately for two feature sets derived from LC1(M) and LC1(P): features selected based on stable sparse weights only, and an expanded set additionally including features with stable loadings. For each pathway and feature set, the number and identity of overlapping inflammation proteins and metabolites are reported alongside pathway-level and joint enrichment p-values.

| **Pathway** | **Feature set** | **Overlapping proteins/genes (n)** | **Overlapping proteins/genes** | **Total pathway proteins/genes** | **Gene enrichment P-value** | **Overlapping metabolites (n)** | **Overlapping metabolites** | **Total pathway metabolites** | **Metabolite enrichment P-value** | **Joint enrichment P-value** |
| --- | --- | --- | --- | --- | --- | --- | --- | --- | --- | --- |
| **Class A/1 (Rhodopsin-like receptors)** | Stable weights | 9 | CXCL6;CXCL1;CXCL10;CCL7;CCL13;CXCL8;CCL19;CCL20;CCL28 | 19 (346) | 0.1110 | 1 | HMDB0000208 | 20 (113) | 0.82200 | 0.31000 |
|  | Stable weights & Stable loadings | 18 | CCL23;CCL2;CXCL6;CXCL11;CXCL5;CXCL1;CX3CL1;CXCL10;CCL3;CXCL9;CCL4;CCL7;CCL11;CCL13;CXCL8;CCL19;CCL20;CCL28 | 19 (346) | 0.0234 | 8 | HMDB0000220;HMDB0000827;HMDB0000208;HMDB0002226;HMDB0000806;HMDB0003229;HMDB0000207;HMDB0002183 | 20 (113) | 0.01410 | 0.00298 |
| **G alpha (q) signalling events** | Stable weights | 1 | MMP1 | 2 (221) | 0.5530 | 0 | NA | 18 (64) | 1.00000 | 0.88100 |
|  | Stable weights & Stable loadings | 2 | MMP1;CCL23 | 2 (221) | 0.5730 | 8 | HMDB0003229;HMDB0002226;HMDB0000827;HMDB0000806;HMDB0000220;HMDB0000207;HMDB0000182;HMDB0002183 | 18 (64) | 0.00654 | 0.02470 |
| **GPCR downstream signalling** | Stable weights | 9 | CXCL6;CXCL1;CXCL10;MMP1;CCL13;CXCL8;CCL19;CCL20;CCL28 | 16 (633) | 0.0318 | 1 | HMDB0000208 | 29 (152) | 0.92400 | 0.13300 |
|  | Stable weights & Stable loadings | 15 | CCL23;CXCL9;CXCL6;CXCL11;CXCL5;CXCL1;CX3CL1;CXCL10;CCL4;MMP1;CCL13;CXCL8;CCL19;CCL20;CCL28 | 16 (633) | 0.0552 | 9 | HMDB0000220;HMDB0000827;HMDB0000208;HMDB0002226;HMDB0000806;HMDB0003229;HMDB0000207;HMDB0000182;HMDB0002183 | 29 (152) | 0.05360 | 0.02020 |
| **GPCR ligand binding** | Stable weights | 9 | CXCL6;CXCL1;CXCL10;CCL7;CCL13;CXCL8;CCL19;CCL20;CCL28 | 19 (476) | 0.1110 | 1 | HMDB0000208 | 27 (168) | 0.90800 | 0.33200 |
|  | Stable weights & Stable loadings | 18 | CCL23;CCL2;CXCL6;CXCL11;CXCL5;CXCL1;CX3CL1;CXCL10;CCL3;CXCL9;CCL4;CCL7;CCL11;CCL13;CXCL8;CCL19;CCL20;CCL28 | 19 (476) | 0.0234 | 9 | HMDB0000220;HMDB0000827;HMDB0000208;HMDB0002226;HMDB0000806;HMDB0003229;HMDB0000207;HMDB0000182;HMDB0002183 | 27 (168) | 0.03360 | 0.00641 |
| **Signaling by GPCR** | Stable weights | 10 | CXCL6;CXCL1;CXCL10;MMP1;CCL7;CCL13;CXCL8;CCL19;CCL20;CCL28 | 20 (706) | 0.0608 | 1 | HMDB0000208 | 30 (190) | 0.93100 | 0.21900 |
|  | Stable weights & Stable loadings | 19 | CCL23;CCL2;CXCL6;CXCL11;CXCL5;CXCL1;CX3CL1;CXCL10;CCL3;CXCL9;CCL4;MMP1;CCL7;CCL11;CCL13;CXCL8;CCL19;CCL20;CCL28 | 20 (706) | 0.0173 | 9 | HMDB0000220;HMDB0000827;HMDB0000208;HMDB0002226;HMDB0000806;HMDB0003229;HMDB0000207;HMDB0000182;HMDB0002183 | 30 (190) | 0.06610 | 0.00891 |

**Notes:** Overlapping proteins/genes: number of submitted proteins overlapping with pathway members. Total pathway proteins/genes: number of pathway members present in the submitted feature list, with total pathway size in parentheses. Overlapping metabolites: number of submitted metabolites overlapping with pathway members. Total pathway metabolites: number of pathway metabolites present in the submitted feature list, with total pathway size in parentheses. Gene enrichment p-value and metabolite enrichment p-value: one-sided hypergeometric test p-values computed separately for proteins and metabolites. Joint enrichment p-value: combined p-value computed by IMPaLA integrating both protein and metabolite enrichment signals simultaneously. Stable weights: features assigned non-zero sparse weights in ≥80% of resampling iterations with consistent directionality. Stable weights and stable loadings: stable weight features additionally supplemented with features exhibiting mean absolute loadings ≥0.3 across resampling iterations. NA indicates no overlapping metabolites were identified for that pathway and feature set. HMDB identifiers are reported for overlapping metabolites.

**Supplementary Table 7**. Baseline characteristics of study population

|  | **Total study population, n (%)** | | **^a^At-diagnosis population, n (%)** | | **^a^Pre-diagnosis population, n (%)** | | **^b^P-value (pre-diagnosis vs. at-diagnosis population)** | **^b^P-value**  **(pre-diagnosis vs. at-diagnosis within Cases)** | **^b^P-value**  **(pre-diagnosis vs. at-diagnosis within Controls)** |
| --- | --- | --- | --- | --- | --- | --- | --- | --- | --- |
|  | **Cases (N=81)** | **Controls (N=81)** | **Cases (N=51)** | **Controls (N=53)** | **Cases (N=30)** | **Controls (N=28)** |  |  |  |
| **Age at sample collection (years)** |  |  |  |  |  |  | 0.521 | 0.116 | 0.261 |
| Mean (SD) | 45.9 (14.9) | 45.6 (15.6) | 43.8 (13.6) | 44.2 (15.0) | 49.5 (16.4) | 48.4 (16.5) |  |  |  |
| Median [IQR] | 43.0 [ 24.0] | 43.0 [ 24.0] | 42.0 [ 22.5] | 42.0 [ 23.0] | 50.5 [ 22.2] | 48.5 [ 21.2] |  |  |  |
| **Sex** |  |  |  |  |  |  | 0.945 | 0.848 | 0.522 |
| Female | 67 (82.7%) | 68 (84.0%) | 43 (84.3%) | 46 (86.8%) | 24 (80.0%) | 22 (78.6%) |  |  |  |
| Male | 14 (17.3%) | 13 (16.0%) | 8 (15.7%) | 7 (13.2%) | 6 (20.0%) | 6 (21.4%) |  |  |  |
| **BMI(kg/m^2^)** |  |  |  |  |  |  | 0.52 | 0.138 | 0.247 |
| Mean (SD) | 28.3 (7.2) | 27.9 (6.7) | 27.4 (7.2) | 27.3 (6.4) | 29.8 (7.0) | 29.1 (7.0) |  |  |  |
| Median [IQR] | 27.1 [ 7.9] | 26.6 [ 7.7] | 25.9 [ 8.1] | 26.2 [ 7.4] | 29.1 [ 9.4] | 27.2 [ 8.5] |  |  |  |
| **Race/Ethnicity** |  |  |  |  |  |  | 0.986 | 0.341 | 0.263 |
| African American | 12 (14.8%) | 12 (14.8%) | 5 (9.8%) | 5 (9.4%) | 7 (23.3%) | 7 (25.0%) |  |  |  |
| East or Southeast Asian | 3 (3.7%) | 3 (3.7%) | 2 (3.9%) | 2 (3.8%) | 1 (3.3%) | 1 (3.6%) |  |  |  |
| European American | 39 (48.1%) | 40 (49.4%) | 28 (54.9%) | 30 (56.6%) | 11 (36.7%) | 10 (35.7%) |  |  |  |
| Hispanic American | 19 (23.5%) | 19 (23.5%) | 12 (23.5%) | 12 (22.6%) | 7 (23.3%) | 7 (25.0%) |  |  |  |
| Other | 8 (9.9%) | 7 (8.6%) | 4 (7.8%) | 4 (7.5%) | 4 (13.3%) | 3 (10.7%) |  |  |  |
| **Storage time (years)** |  |  |  |  |  |  | 0.303 | 0.037 | 0.071 |
| Mean (SD) | 9.2 (3.3) | 9.2 (3.2) | 8.7 (3.4) | 8.7 (3.4) | 10.2 (2.8) | 10.0 (2.8) |  |  |  |
| Median [IQR] | 10.0 [ 3.0] | 10.0 [ 2.0] | 10.0 [ 6.0] | 10.0 [ 6.0] | 11.0 [ 3.0] | 11.0 [ 3.0] |  |  |  |
| **^c^Time between plasma sample collection and thyroid cancer diagnosis / reference time (years)** |  |  |  |  |  |  | <0.001 | <0.001 | <0.001 |
| Mean (SD) | 1.5 (2.4) | 1.4 (2.3) | 0.1 (0.2) | 0.1 (0.2) | 4.0 (2.3) | 4.0 (2.3) |  |  |  |
| Median [IQR] | 0.1 [2.0] | 0.0 [1.9] | 0.0 [0.0] | 0.0[0.0] | 3.7 [3.9] | 3.7 [3.9] |  |  |  |

SD: standard deviation. IQR: interquartile range.

^a^Pre-diagnosis indicates samples collected > 1 year before diagnosis; at-diagnosis indicates samples collected within 1 year before diagnosis. The thyroid cancer cases and matched controls were drawn from BioMe, a medical record-linked biobank at the Icahn School of Medicine at Mount Sinai. Controls were individually matched to cases by sex, age at sample collection, race/ethnicity, BMI, and sample collection year^1,7^.

^b^In comparing the pre-diagnosis study population to the at-diagnosis study population, we applied the Independent T-test for continuous variables and Fisher’s Exact Test for categorical variable

^c^For cases: time between plasma sample collection and cancer diagnosis. For controls: reference time assigned by study design (equals the matched case's time to diagnosis^1,7^.

**Supplementary Table 8: Metrics used to assess component-level and feature-level stability**

Stability was assessed across resampling at the latent component (LC) and feature levels using complementary metrics of reproducibility, selection consistency, and association strength. In-sample data were used to evaluate feature selection and weights, and out-of-sample data to assess projected scores and loadings.

| **Aspect** | **Variables** | **Dataset** | **Stability Metrics** | **Evaluation Objective** |
| --- | --- | --- | --- | --- |
| Latent Component | LC score  (sample scores) | Out-of-sample | Spearman’s rho (score rank) | Reproducibility of sample projections across resampling |
|  | LC Feature Selection | In-sample | Jaccard index^8^  Nogueira Index^9,10^ | Consistency of selected feature subsets |
|  | LC Sparse Weight Vector | In-sample | Spearman’s  rho (weight rank) | Preservation of feature ranking across resampling |
| Feature | Feature Weights | In-Sample | Selection Frequency (SF)  Sign Consistency (SC) | Stability of feature selection;  Directional stability of feature weights |
|  | Feature Loadings  (correlation) | Out-of-Sample | Sign Consistency | Directional consistency of feature–LC correlation;  Strength of feature–LC correlation |

**Note:** Stability metrics are reported as continuous measures and summarized across resampling. Higher values indicate greater reproducibility or consistency. For interpretability, commonly used reference ranges (e.g., 0.5 for Spearman’s rho, 0.8 for sign consistency, and 0.3 for loading magnitude) may be considered as descriptive guidelines, but are not treated as formal thresholds. The Nogueira index accounts for chance agreement under sparsity, whereas the Jaccard index does not.

**Reference**

1.	Van Gerwen, M. *et al.* Per- and polyfluoroalkyl substances (PFAS) exposure and thyroid cancer risk. *eBioMedicine* **97**, 104831 (2023).

2.	India Aldana, S. *et al.* Pregnancy as a Susceptible Period to Ambient Air Pollution Exposure on the Maternal Postpartum Metabolome. *Environ. Sci. Technol.* **59**, 6400–6413 (2025).

3.	Schymanski, E. L. *et al.* Identifying Small Molecules via High Resolution Mass Spectrometry: Communicating Confidence. *Environ. Sci. Technol.* **48**, 2097–2098 (2014).

4.	Djoumbou Feunang, Y. *et al.* ClassyFire: automated chemical classification with a comprehensive, computable taxonomy. *J. Cheminformatics* **8**, 61 (2016).

5.	Kirwan, J. A., Broadhurst, D. I., Davidson, R. L. & Viant, M. R. Characterising and correcting batch variation in an automated direct infusion mass spectrometry (DIMS) metabolomics workflow. *Anal. Bioanal. Chem.* **405**, 5147–5157 (2013).

6.	Stekhoven, D. J. & Bühlmann, P. MissForest—non-parametric missing value imputation for mixed-type data. *Bioinformatics* **28**, 112–118 (2012).

7.	Joseph, G. *et al.* Inflammatory proteins in pre‐diagnosis versus at‐diagnosis samples associated with differentiated thyroid cancer. *Int. J. Cancer* **158**, 2072–2082 (2026).

8.	Hédou, J. *et al.* Discovery of sparse, reliable omic biomarkers with Stabl. *Nat. Biotechnol.* **42**, 1581–1593 (2024).

9.	Meinshausen, N. & Bühlmann, P. Stability Selection. *J. R. Stat. Soc. Ser. B Stat. Methodol.* **72**, 417–473 (2010).

10.	Nogueira, S., Sechidis, K. & Brown, G. On the Stability of Feature Selection Algorithms.

11.	Karnovsky, A. *et al.* Metscape 2 bioinformatics tool for the analysis and visualization of metabolomics and gene expression data. *Bioinformatics* **28**, 373–380 (2012).

12.	Assenov, Y., Ramírez, F., Schelhorn, S.-E., Lengauer, T. & Albrecht, M. Computing topological parameters of biological networks. *Bioinformatics* **24**, 282–284 (2008).

13.	Shannon, P. *et al.* Cytoscape: A Software Environment for Integrated Models of Biomolecular Interaction Networks. *Genome Res.* **13**, 2498–2504 (2003).
